# Long read metagenomics-based precise tracking of bacterial strains and genomic changes after fecal microbiota transplantation

**DOI:** 10.1101/2024.09.30.615906

**Authors:** Yu Fan, Mi Ni, Varun Aggarwala, Edward A. Mead, Magdalena Ksiezarek, Lei Cao, Michael A. Kamm, Thomas J. Borody, Sudarshan Paramsothy, Nadeem O. Kaakoush, Ari Grinspan, Jeremiah J. Faith, Gang Fang

## Abstract

Fecal microbiota transplantation (FMT) has revolutionized the treatment of recurrent *Clostridioides difficile* infection (rCDI) and is being evaluated across other diseases. Accurate tracking of bacterial strains that stably engraft in recipients is critical for understanding the determinants of strain engraftment, evaluating their correlation with clinical outcomes, and guiding the development of therapeutic bacterial consortia. While short-read sequencing has advanced FMT research, it faces challenges in strain-level *de novo* metagenomic assembly. In this study, we described a novel framework, *LongTrack*, which uses long-read metagenomic assemblies and rigorous informatics tailored for FMT strain tracking. We highlighted LongTrack’s advantage over short-read approaches especially when multiple strains co-exist in the same sample. We showed LongTrack uncovered hundreds of engrafted strains across six FMT cases of rCDI and inflammatory bowel disease patients. Furthermore, long reads also allowed us to assess the genomic and epigenomic stability of engrafted strains during the 5-year follow-ups, revealing structural variations that may be associated with strain adaptation in a new host environment. Combined, our study advocates the use of long-read metagenomics and LongTrack to enhance strain tracking in future FMT studies, paving the way for the development of more effective defined biotherapeutic as an alternative to FMT.

## Main

Fecal microbiota transplantation (FMT) has emerged as a new therapeutic strategy, transforming the treatment paradigms for patients with recurrent *Clostridioides difficile* infection (rCDI)^1–3^. By transferring gut microbiota from a healthy donor into a patient, FMT effectively restores the patient’s gut microbial ecology^4,5^. Potential applications of FMT extend beyond rCDI onto treating a broader array of diseases^6,7^, such as the treatment of inflammatory bowel disease (IBD) ^8–10^ and cancer immunotherapy^11–13^.

The FMT procedure operates on the principle of transferring gut microbiota, which constitutes diverse bacterial strains, the functional impact units of the gut microbiota^14–16^, from a healthy donor into a recipient^4,17^. Precise and comprehensive tracking of these strains from transplantation to engraftment within the recipient’s gut is essential for understanding factors that facilitate or hinder colonization^5,17–19^, providing vital information on how specific strains correlates with clinical outcomes^18,20^, which can guide the development of defined bacterial consortiums, composed of strains with beneficial effects, for safer therapeutic applications^19–21^.

High throughput metagenomic shotgun sequencing studies have provided valuable insights into strain engraftment in FMT in various diseases^18,20–25^ primarily through short-read sequencing to detect single nucleotide polymorphisms (SNPs) and/or gene clusters that can track strains in the recipient after FMT^22–24,26–28^. However, a recent study, using genome sequences from extensive bacterial isolates collection^20^ for strain tracking, showed that short-read approaches face challenges when multiple strains co-exist in a single microbiome sample^29–34,35^. Once individual isolate genomes from donor and pre-FMT recipient samples are assembled with high-purity and high-completeness, reliable tracking can be achieved using each strain’s unique k-mers in short-read metagenomic data^20^. However, standard culturing methods often miss certain bacterial taxa due to the selectivity of culture media and/or conditions^36^, and pose challenges for scaling up large FMT cohorts. Long-read sequencing empowers *de novo* genome assembly of complete, contiguous, strain-level resolution metagenome-assembled genomes (MAGs) at strain resolution^32,37–39^, greatly complementing culture and short-read approaches^40–42^. With this rationale, we developed a new framework, *LongTrack*, that uses rigorous informatics tailored to reliably use long-read MAGs for strain tracking (**Fig. 1**). Building upon systematically cultured bacterial strains from FMT donors and recipients for rCDI with 5-year follow-ups^20^ (**Fig. 1A**), this strain collection served as a ground truth for LongTrack evaluation (**Fig. 1B**) and showed that LongTrack outperforms short-read based methods, especially in distinguishing co-existing bacterial strains within a microbiome sample (**Fig. 1C**). Additionally, taking the advantages of long-read sequencing in strain-level read mapping^40,42^ and direct detection of bacterial DNA methylation^43–47^, we identified structural variations (SVs) and epigenetic changes within strains as they transferred from donors to post-FMT recipients during follow-up (**Fig. 1D**).

**Figure 1.**
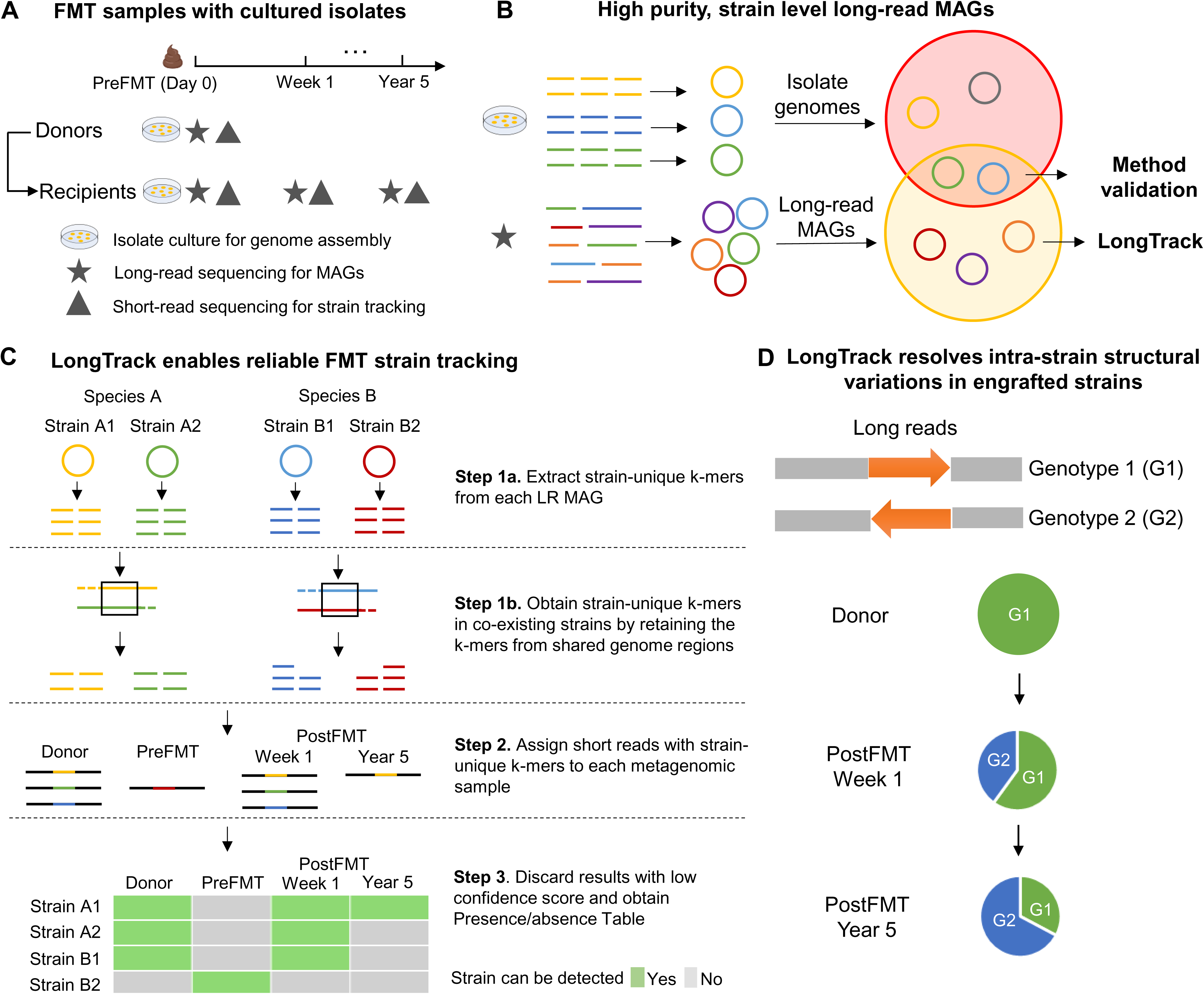
Description of FMT samples, matched bacteria isolates and the core design of LongTrack for reliable FMT strain tracking and detecting intra-strain structural variations in engrafted strains. **A**, Stool samples were collected from three FMT donors and four FMT recipients at multiple time points over five years. Extensive bacterial culture, isolate genome assemblies, and short read-metagenomic sequencing have been performed in the previous study by Aggarwala et al and shown along with FMT samples. Long-read metagenomic sequencing data were newly generated for this study. **B**, Long-read data from donors’ and recipients’ samples before FMT were used to construct *de novo* metagenome-assembled genomes (long-read MAGs). The overlapping isolate genomes enable a systematic evaluation of the reliability of long-read MAGs, which serves as the foundation of LongTrack. **C**, Schematics of the core design of LongTrack for FMT strain tracking based on strain-level long-read (LR) MAGs. Strain-unique k-mers (k=31) were identified for each strain-level LR MAG from donor or pre-FMT recipient as the markers by eliminating common k-mers present in public database, or the other MAGs in the same FMT donor and recipient (**Methods**). To ensure specificity, strain-unique k-mers are only selected from shared regions across multiple strain-level LR MAGs within the same species. The presence of these final unique k-mers of a specific MAG (above a confidence score, **Methods**) in a post-FMT recipient’s short-read metagenomic data indicates that the corresponding MAG is present (green color) in the post-FMT recipient sample. **D**, Long-read metagenomic sequencing data were also utilized for strain-specific detection of structure variations (SVs) that differ between donor and post-FMT samples. The references for putative genotypes were created and used to map and quantify specific SVs in each sample, allowing tracking of SV changes between hosts and/or along time. The long-read approach has unique advantage in strain-level read mapping that allows reliably detection of intra-strain variations, which are challenging for short reads due to their limitations in accurately mapping reads to specific strains.

## Results

### High purity, strain-level long-read MAGs serve as a solid foundation for LongTrack

The success of strain tracking in FMT depends on obtaining strain-resolved and high purity genomes from both the donor and the recipient before FMT, allowing differentiation among co-existing strains of the same species.

Bacterial isolation is an effective method for achieving this, as sequencing individual colonies yield discrete genomes with high purity and completeness. With this rationale, we built on the collection of 1000+ bacteria isolates with individually assembled genomes as ground truth^20^ to evaluate the long-read MAGs, and their accuracy in FMT strain tracking. We selected four FMT cases of rCDI (3 donors and 4 recipients) from which, 184 unique strains (105 species) were isolated with individually assembled genomes^20^ (**Extended Data Fig. 1**). We generated new long-read sequencing data for 15 samples, and deeper short-read for two samples to enable a systematic evaluation along with LongTrack.

To construct strain-level long-read MAGs, we employed a pipeline tailored for long-read sequencing data, involving metagenomic assembly, haplotype phasing, and strain-level refinement (**Extended Data Fig. 2, Methods**). From four cases, we obtained a total of 460 long-read MAGs (460 strains across 312 species), including 60 strains shared with cultured isolates (**Fig. 2A**). As Donor 1 (D1) had the most bacterial isolates (136 unique strains), we focused on D1 to demonstrate the advantage of long reads for *de novo* assembly of MAGs compared to short reads. We obtained 135 short-read MAGs from D1 (**Fig. 2B**) and 20 short-read MAGs from R1 using metaSPAdes^48^, which demonstrates better performance compared to MEGAHIT^49^ (**Supp Fig. S1).** Contamination analysis with CheckM^50^ for 460 long-read MAGs (from all 4 cases) and 155 short-read MAGs (from D1 and R1) revealed lower contamination and higher contig N50 values in long-read MAGs than short-read MAGs (**Fig. 2C-D**). Specifically, for D1, the assemblies yielded 124 long-read MAGs and 135 short-read MAGs with comparable completeness (78.2% vs 78.6 %, **Extended Data Fig. 3A**). Although short-read data had the advantage in sequencing depth (47 Gb vs. 23 Gb), long-read MAGs still consistently exhibit significantly lower contamination rates (1.24% vs 2.11%, **Extended Data Fig. 3B**) and larger N50 values (627.5 kb vs 56.9 kb, **Extended Data Fig. 3C**). These findings are consistent with previous studies that demonstrated the advantage of long-read sequencing for *de novo* metagenomic assembly across many microbiome samples^30,37–40,51,52^.

**Figure 2.**
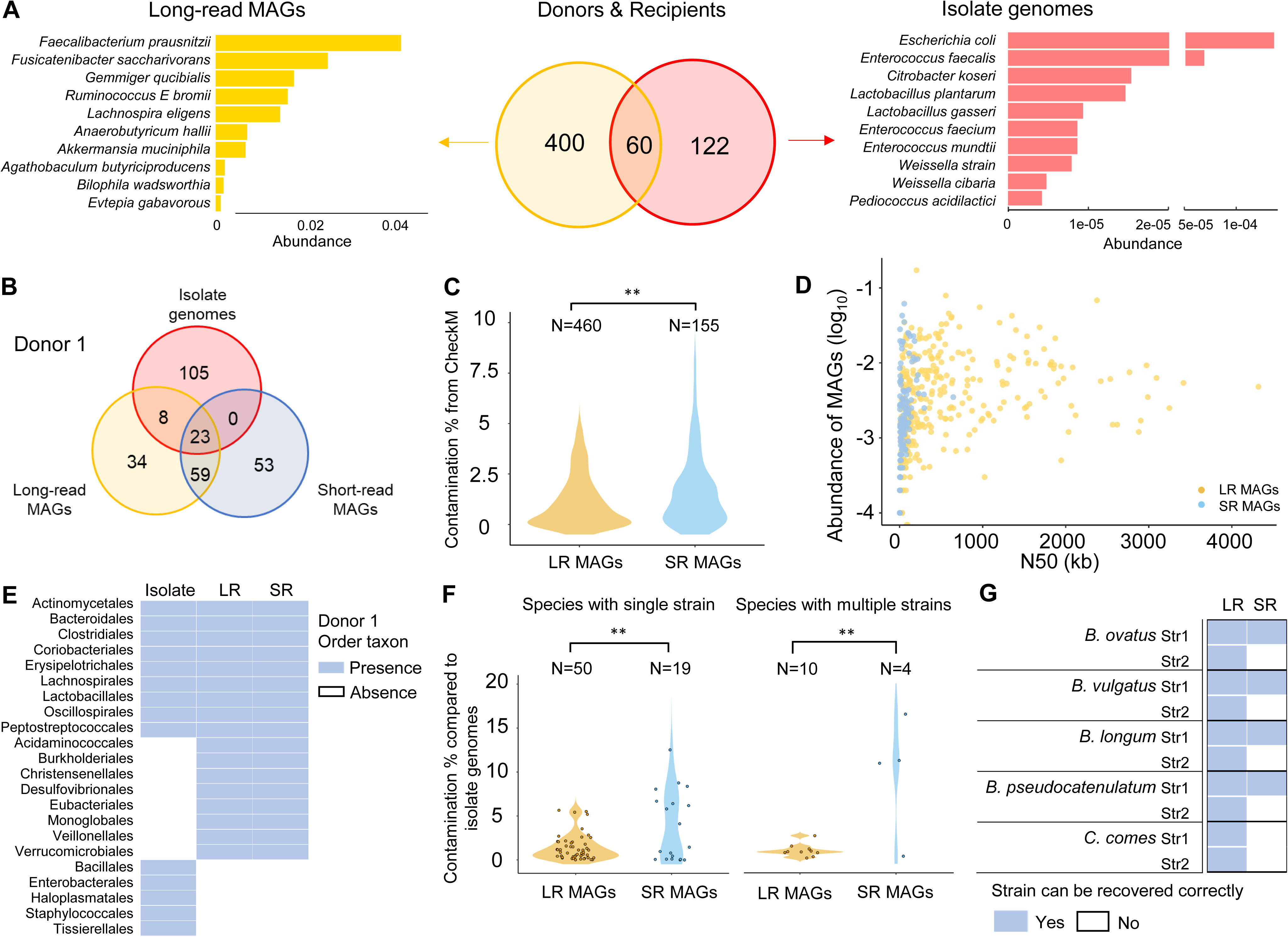
High-purity, strain-level long-read MAGs serve as a solid foundation for LongTrack. **A**, The Venn diagram displays the number of strains that were recovered by long-read (LR) MAGs (yellow color) or isolate genomes from bacterial culture (red color) from 3 donors and 4 recipients, with the left bar plot illustrating examples of species with MAGs with modest-to-high relative abundance but were missed by culture, and with right bar plot illustrating examples of isolates that were cultured but not recovered by long-read sequencing because of their very low abundances. **B**, The Venn diagram displays the number of strains that can be recovered by LR MAGs (yellow color), short-read (SR) MAGs (blue) or isolate genomes (red) in Donor 1 (D1). LR MAGs include all the 23 strains recovered by SR MAGs, along with 8 additional strains not assembled by short reads. **C**, Contamination levels of 460 LR MAGs (from Donor 1-3 and Recipient 1-4) and 135 SR MAGs (from Donor 1, D1, for which we generated ultra-deep data) assessed using CheckM, showing significantly lower contamination in LR MAGs (p < 0.01, **, Wilcoxon test). **D**, Distributions of relative abundance (log scale) and contig N50 (kb) for LR MAGs (from Donor 1-3 and Recipient 1-4) and SR MAGs (from D1). **E,** Order-level taxonomic comparison of LR MAGs, SR MAGs, and isolate genomes from D1. **F**, Contamination level for 60 LR MAGs (from Donor 1-3 and Recipient 1-4) and 23 SR MAGs (from D1), evaluated across various species with a single strain or multiple strains. LR MAGs consistently show lower contamination (p < 0.01, **, Permutation test). Contamination levels were assessed by mapping long-read and short-read MAGs to their corresponding isolate genomes and calculating the percentage of unaligned bases as a measure of contamination. **G**, LR sequencing have a unique advantage to resolve multiple co-existing strains from the same species than SR sequencing.

In our in-depth evaluation, we compared these MAGs to the isolate genomes. For D1, 31 and 23 isolate genomes were shared with long-read MAGs and short-read MAGs, respectively (**Fig. 2B**). and long-read MAGs include all 23 MAGs recovered by short-read MAGs along with 8 additional strains (**Fig. 2B**). Both long-read and short-read MAGs spanned the same 17 taxonomic orders, with 8 detected only by MAGs (**Fig. 2E**). Additionally, all MAGs overlapping with isolate genomes, including 60 long-read MAGs (across 4 cases) and 23 short-read MAGs (from D1), were used to evaluate the reliability of long-read MAGs in comparison with short-read MAGs. First, we assessed their contamination levels by mapping MAGs to their corresponding isolate genomes (**Fig. 2F, Extended Data Fig. 4A**), which provide more reliable insights than estimates by CheckM^50^ (**Extended Data Fig. 4B)**. The contamination levels were lower for long-read MAGs (1.34%) compared to short-read MAGs (4.77%), for both species with a single and co-existing strains (**Fig. 2F**, **Extended Data Fig. 4C-D**). We further evaluated the k-mer similarity between MAGs and their corresponding isolate genomes, which consistently indicated long-read MAGs exhibit a higher similarity (**Extended Data Fig. 4E**). Importantly, long-read sequencing effectively recovered the same number of strains as bacterial culture from the same donor sample while these co-existing strains were not completely recovered in the short-read MAGs (**Fig. 2G**). We also identified 11 plasmids from the 60 shared isolate genomes of which 7 plasmids, including one circular, were recovered from long-read sequencing. However, none of the 7 plasmids were correctly binned with their matching chromosomes, highlighting that plasmid binning in microbiome settings remain a challenge and need to be assisted by more specialized methods such as Hi-C^53^. While the shared strains demonstrate the reliability of long-read MAGs, the inherent methodological differences between metagenomics and bacterial isolation remain. Specifically, long-read MAGs, favor strains with modest-to-high relative abundance, such as *Akkermansia muciniphila* (0.63%) and *Faecalibacterium prausnitzii* (3.94%), while bacterial culture favors strains selected by culture media rather than abundance, such as *Escherichia coli* with abundance below 0.01% (**Fig. 2A**). In D1, 37.1% and 65.9% metagenomic reads can be mapped to the isolate genomes and long-read MAGs (**Methods**), respectively, with 76% mapping to the merged reference (isolate genomes and long-read MAGs) (**Supp Fig. S2**). Interestingly, the 31 shared strains accounted for 30.7% metagenomic reads, because of their relatively high abundance, highlighting the complementarity between long-read MAGs and bacterial culture.

### LongTrack enables reliable use of Long-read MAGs for precise FMT strain tracking

Once individual isolate genomes are assembled, strain tracking can be reliably performed using unique k-mers (n=31) selected from isolate genomes and short-read sequencing of post-FMT samples, providing a robust ground truth for evaluating LongTrack^20^. This k-mer based strategy (Strainer) has been demonstrated^20^ with superior performance compared to several SNP-based approaches^20^ including inStrain^27^, StrainFinder^23^, and ConStrains^22^, particularly for modest-to-low abundance strains and multiple co-existing strains. Our independent evaluation of LongTrack confirmed that the k-mer based approach is more sensitive than the SNP-based method (StrainFinder^23^), which tends to miss modest-to-low abundance strains (**Extended Data Fig. 5**).

However, Strainer is not compatible with long-read MAGs that often comprise incomplete MAGs (taxa with modest-to-low abundance) which can lead to false strain tracking results) Therefore, to reliably use long-read MAGs for FMT strain tracking, we developed LongTrack, where for similar co-existing strains in one fecal sample, or two similar strains each in the donor sample and recipient pre-FMT samples, we selected unique k-mers only from their common genomic regions (**Fig. 1C**). This adaptation avoids the selection of unreliable k-mers that appear to be unique to a strain but are not, due to the incompleteness of long-read MAGs, highlighting a key difference between high purity but incomplete long-read MAGs and the complete genomes of bacterial isolates^20^ (**Methods**). We compared the number of unique k-mers used for strain tracking in multiple co-existing strains, using the original Strainer with high completeness isolate genomes versus LongTrack with high-purity long-read MAGs (**Fig. 3A**). Although LongTrack generally kept fewer unique k-mers compared to Strainer, it still retained enough unique k-mers for reliable strain tracking in post-FMT samples (**Fig 3B**). In addition, we compared the sensitivity of LongTrack and culture-based strain tracking using the union of all strains detected by long-read MAGs and isolate genomes as the reference set. Specifically, when assessing sensitivity across varying abundance ranges, LongTrack demonstrated 100% sensitivity for strains with relative abundance >0.01 and maintained a high sensitivity of 83% for strains with relative abundance between 0.001 and 0.01, while culture-based strain tracking missed many taxa in these ranges. However, for ultra-low abundance strains (<1e–4), LongTrack’s sensitivity dropped sharply, while culture-based approach achieved >98% sensitivity (**Extended Data Fig. 6**). These results illustrate that LongTrack effectively captures strains with moderate to high abundance, while culture-based methods have a clear advantage in detecting very low-abundance taxa, due to the limited sequencing depth inherent in metagenomic approaches. This evaluation highlights the complementary nature of the two approaches for FMT strain tracking.

**Figure 3.**
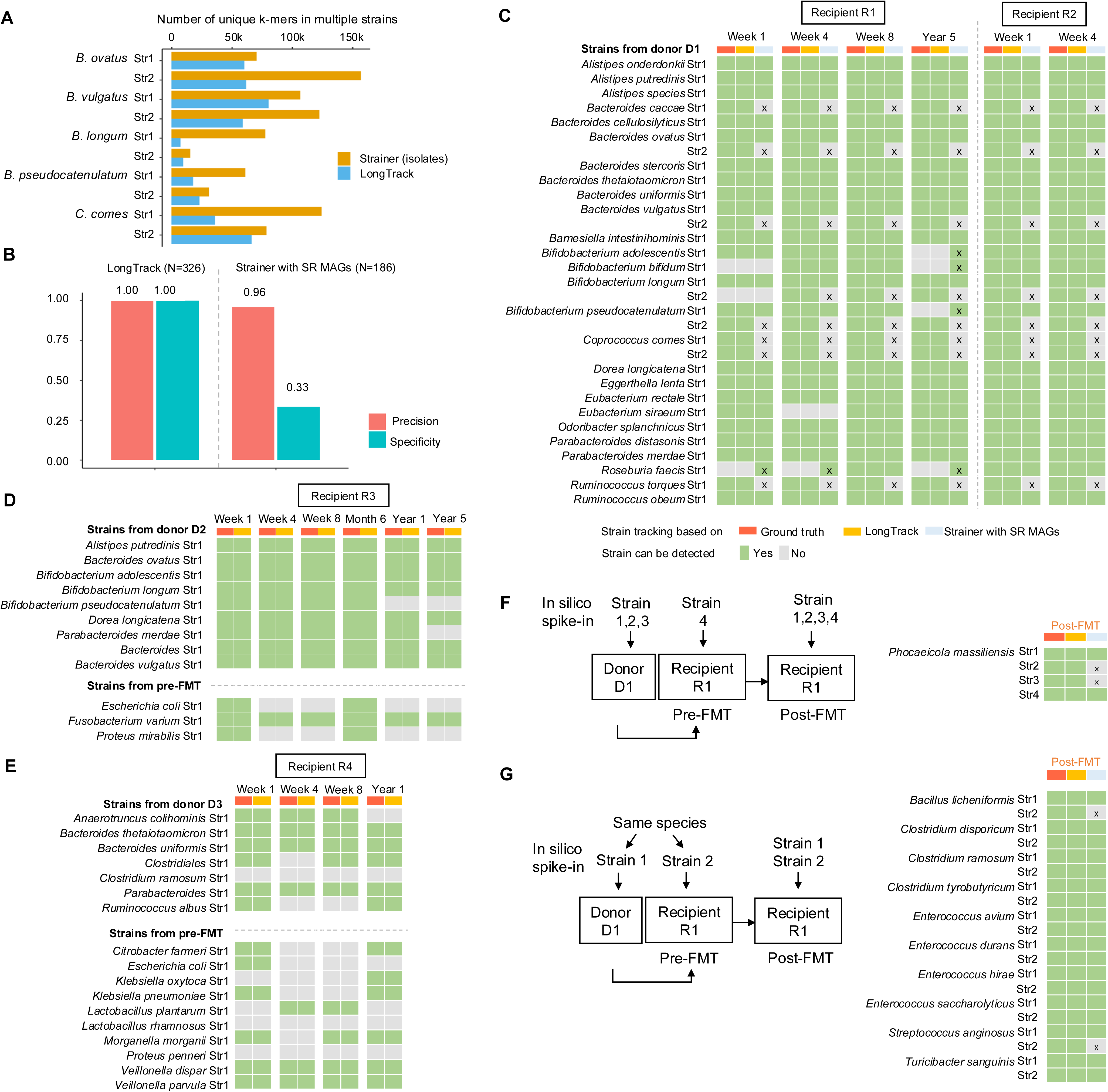
Evaluation of LongTrack for FMT strain tracking. **A**, Numbers of unique k-mers used for strain tracking in cases of multiple co-existing strains by the original Strainer method (with complete genomes assembled for isolates) and the LongTrack method (with long-read MAGs). Although some k-mers from shared regions are filtered out by LongTrack, enough unique k-mers remain for reliable strain tracking in post-FMT samples. **B**, Precision and specificity of strain tracking using LongTrack with long-read MAGs (Donor 1-3 and Recipient 1-4) versus Strainer with short-read MAGs (Donor 1). Across 3 donors and 4 recipients with multiple fecal sampling time points (326 unique strain presence/absence events), LongTrack with long-read MAGs achieved high precision (1.00) and specificity (1.00). In contrast, Strainer with short-read MAGs (n = 186) showed lower precision (0.96) and specificity (0.33). **C**, Presence (green) or absence (gray) of strains in post-FMT recipients determined by strain-specific unique k-mers selected from the ground truth (Strainer with isolate genomes), LongTrack with long-read MAGs, or Strainer with short-read MAGs. Discrepancies with the ground truth are indicated by an "X" marker. **(C-E).** The evaluation of strain tracking was performed for four FMT cases of rCDI: D1->R1 (**C**), D1->R2 (**C**), D2->R3 (**D**), and D3->R4 (**E**). **F**, Evaluation of LongTrack for FMT strain tracking using *in silico* spike-in data that represent the co-existence of multiple strains of the same species in the donor sample. Specifically, we spiked-in three strains of *Phocaeicola massiliensis* (Str 1-3) to D1, and a fourth strain of *P. massiliensis* (Str 4) to R1 pre-FMT. All four strains were added to post-FMT recipient R1. **G**, Evaluation of LongTrack for FMT strain tracking using *in silico* spike-in data that represent cases when there are high-similarity strains present in between the donor and pre-FMT samples. Specifically, we spiked-in ten pairs of strains, each with high similarity (ANI>99.5%), from ten different species, one strain to D1 and the other to pre-FMT recipient R1, similarly for all the ten pairs. All twenty strains were added to post-FMT recipient R1.

We further compared precision and specificity of strain tracking based on LongTrack with long-read MAGs (across 4 cases) versus Strainer with short-read MAGs (D1), with multiple fecal sampling time points each. LongTrack achieved great precision (1.00) and specificity (1.00) (**Fig. 3B**) when using shared isolate genomes as the ground truth^20^, while Strainer with short-read MAGs had lower precision (0.96) and much lower specificity (0.33) (**Fig. 3B**), likely due to the false-positive tracking and missed the co-existing strains (**Fig. 3C**). Building upon the two well-characterized FMT cases, D1->R1 and D1-> R2, we found 100% consistency of strain tracking results between LongTrack with long-read MAGs and ground truth for all 31 shared strains (**Fig. 3C**). In contrast, Strainer with short-read MAGs correctly track 23 of the 31 strains (74%) but failed to track the other 8 strains in post-FMT samples as it was not able to resolve the multiple strains co-existing within the same donor sample. In addition, we also observed that Strainer with short-read MAGs produced six false positives due to high contamination. For instance, the short-read MAGs *Bifidobacterium adolescentis* has contamination levels of 8.76%, with 28% of the contamination originating from *Bacteroides vulgatus* within the same sample (**Supp Fig. S3),** leading to unreliable k-mers and the false positive tracking results (**Fig. 3C**). We applied LongTrack to two additional rCDI FMT cases: D2->R3 and D3->R4 and also found high precision (**Fig. 3D-E**). A noteworthy observation is that long-read MAG completeness does not significantly affect tracking accuracy. We performed additional evaluations with two mock FMT scenarios with *in silico* spike-in (**Methods**). In the first evaluation, we spiked-in three strains of *Phocaeicola massiliensis* (Str 1-3) to D1, and a fourth strain of *P. massiliensis* (Str 4) to pre-FMT R1; all four were present in post-FMT R1 (**Fig. 3F**). LongTrack recovered all four strains and accurately tracked them in post-FMT R1, while short-read MAGs only recovered two of the four strains (**Fig. 3F**). In the second evaluation, we spiked-in ten pairs of strains, each with high similarity (ANI>99.5%), from ten different species, to D1 and pre-FMT R1. All twenty strains were added to post-FMT R1 (**Fig. 3G**). LongTrack correctly tracked all strains in post-FMT recipients, while Strainer with short-read MAGs missed two (**Fig. 3G**).

### LongTrack helps better understand strain engraftment in rCDI and IBD patients

Building on the reliability of LongTrack-based FMT strain tracking as evaluated, we scale up strain tracking to more comprehensively understand strain engraftment in FMT. Across all the four cases of rCDI FMT, long-read sequencing recovered 396 MAGs with diverse taxonomy distribution (**Fig. 4A**) that were not captured by the previous culture-based study^20^.

**Figure 4.**
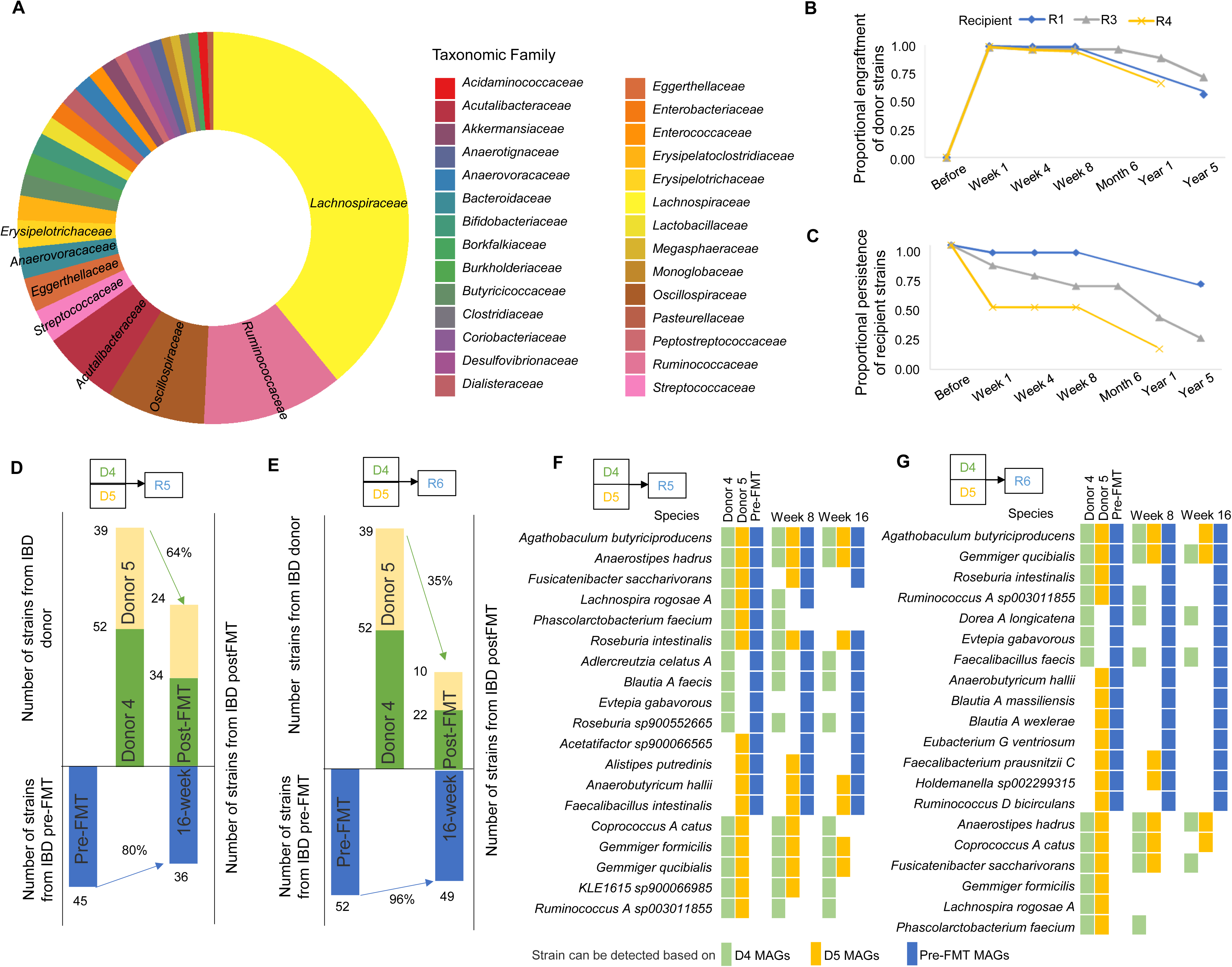
LongTrack uncovered hundreds of engrafted strains across six FMT cases of rCDI and IBD patients. **A**, Diverse taxonomy distribution of long-read MAGs that are not cultured in the previous study by Aggarwala *et al*. **B**, Proportional engraftment of donor strains (PEDS) in individual post-FMT recipients at various time points of the 5-year follow up. **C**, Proportional persistence of recipient strains (PPRS) in individual post-FMT recipients at various time points of the 5-year follow up. **D** & **E**, PEDS and PPRS analysis for the two FMT cases for IBD patients R5 (**D**) and R6 (**E**), where three colors are used to represent the strains from the two Donors (D4 and D5) or the recipient. (**F** & **G**) The presence and absence of co-existing strains in the two IBD patients (R5, **F**, R6, **G**) after FMT that originated from the two Donors (D4 and D5) or the recipient.

We performed LongTrack to measure the engraftment of donor strains in the recipients, focusing on uncultured strains from long read. We used *proportional engraftment of donor strains* (PEDS) to summarize strain engraftment up to five years after FMT. We found consistent and stable engraftment of donor strains (**Fig. 4B**), with PEDS values comparable to cultured strains (**Extended Data Fig. 7A**). In these individuals, we report an average PEDS of 98% of uncultured strains in the first week, which stabilized ∼96% at 8 weeks and remained 63% even 5 years later. This demonstrates that FMT can lead to a stable long-term engraftment of donor‘s uncultured strains in rCDI patients. We also monitored the persistence of recipient strains post-FMT using *proportional persistence of recipient strains (PPRS)*. We observed a major reduction (47%) in PPRS of recipient strains post-FMT, both for cultured and uncultured strains, indicating a major shift in the gut microbiota following FMT (**Fig. 4C, Extended Data Fig. 7B**). The absence of *C. difficile* in post-FMT R1 is noteworthy, suggesting that the transplantation led to a more balanced and health-promoting gut microbiota (**Extended Data Fig. 8A**).

We examined the individual trajectories of long-read MAGs recovered strains for donor-recipient pairs (**Extended Data Fig. 8**) and found several strains that stably engrafted in the recipient after five years potentially linked to human health and diseases. One notable example, *A. muciniphila*, present in D1 and D2, is a mucin-degrading bacterium linked to various health benefits and improved treatment outcomes^54–59^. It helps prevent *C. difficile* overgrowth through the modulating gut microbiota and metabolites^32^, supports gut barrier integrity and immune response^33^. *A. muciniphila* strains from D1 and D2 demonstrated stable engraftment in post-FMT recipients, with persistence observed for up to five years (**Extended Data Fig. 8A-B**). *F. prausnitzii* is another species well resolved exclusively by long-read MAGs. and related to human health through short chain fatty acids production^61–66^ and, a potential role in maintaining gut health and preventing IBD^62,63,67^. *F. prausnitzii* strain from D1 was detected in 8-week post-FMT R1 but not detected 5-year post-FMT R1 (**Extended Data Fig. 8A**).

Next, we performed LongTrack to expand the tracking analysis to strains from post-FMT samples and reversely tracked them in earlier time points (**Methods**). This approach captures strains that were low-abundance in donor or pre-FMT but increased in post-FMT samples, allowing them to be assembled by long reads. Indeed, this design helped us identify 35 additional strains from the three FMT cases (**Extended Data Fig. 9A-C**). All these cases have clear evidence that they were from either the donor sample or pre-FMT samples but were not recovered by long-read sequencing due to low abundance. Interestingly, we also observed novel strains recovered from post-FMT recipients, but no evidence supported their presence in any of the earlier times, neither donor nor pre-FMT (**Extended Data Fig. 9D-F**). For example, *Acidaminococcus intestini,* present only in the 5-year post-FMT R1 (**Extended Data Fig. 9D**), was likely acquired from the environment sometime between week 8 and year 5, unless its abundance was below detections limit in all the earlier time points.

We further applied the LongTrack to two FMT cases for the treatment of IBD^8,68^, which represents pre-FMT recipients with more complex microbiome than rCDI patients. The two IBD patients (R5 and R6) each received fecal samples from six donors (D4 - D9) (**Extended Data Fig. 10**), as multiple donors are often used in the FMT treatment of IBD ^8,68^. This greater complexity of FMT provided a great opportunity to demonstrate the power of LongTrack for strain tracking. We conducted long-read sequencing on two of the donors (D4 and D5) and the two pre-FMT recipients and recovered 188 MAGs. Out of these strains, 64% and 35% strains from the two donors successfully engrafted in the two 16-week post-FMT recipients, respectively (**Figs. 4D-E**). We also estimated PPRS and observed that 80% and 96% of strains from pre-FMT recipients were present in 16-week post-FMT recipients (**Fig. 4D-E**). Compared to the FMT cases for rCDI at a similar time point (8 weeks), we observed lower PEDS and higher PPRS in the FMT cases for IBD, which is consistent with the expectation that the gut microbiome of IBD patients is more diverse and resilient upon FMT introduction of donor strains^8^. We observed 19 and 20 coexisting strains (from donors and pre-FMT recipients) in the two IBD post-FMT recipients (**Fig. 4F-G**). Notably, while nearly all strains from the pre-FMT recipients remained stable for 16-week post-FMT, some donor strains were not detectable, suggesting endogenous strains in IBD patients are more resilient in the same host after FMT.

### Long-read sequencing of post-FMT samples uncovered genomic structural variations within engrafted strains

It is crucial to understand how donor strains adapt to the recipient’s microbiome environment, which can drive genomic variations - that not been extensively studied^69–71^. Long-read sequencing has great advantages in complex SV detection and direct detection of DNA methylation^43–47,72^, enabling to assess the stability and dynamics of SVs and DNA methylation in individual strains across donor and post-FMT

A systematic analysis of long-read sequencing data of the post-FMT recipients across various time points identified 38 SVs, including 6 deletions, 20 insertions, and 12 inversions, as determined by stringent filtering criteria (**Methods**). Comparison of read alignments of SV regions showed that both long-read sequencing methods provided clear and consistent flanking sequence alignments (**Fig. 5A**), illustrating they were from the same strain, whereas short-read sequencing plots displayed several mismatches, making it challenging to reliably attribute the sequences to the same strain (**Fig. 5A**). Out of the 38 SVs, 12 were identified in four strains that we were able to isolate from both donor and post-FMT recipient samples, allowing us to validate these SVs. Long-read sequencing of these newly cultured isolates confirmed paired donor and post-FMT recipient isolates are indeed the same strain, with high genome-wide ANI (>99.9%). Among the 12 SVs, 11 alternative SV genotypes (91.67%) indeed exist in the same strain as validated by long-read sequencing of the isolates from donor and post-FMT samples, demonstrating the high reliability of SV detection within the same strain using long-read sequencing (**Fig. 5B, Supp Fig. S4**).

**Figure 5.**
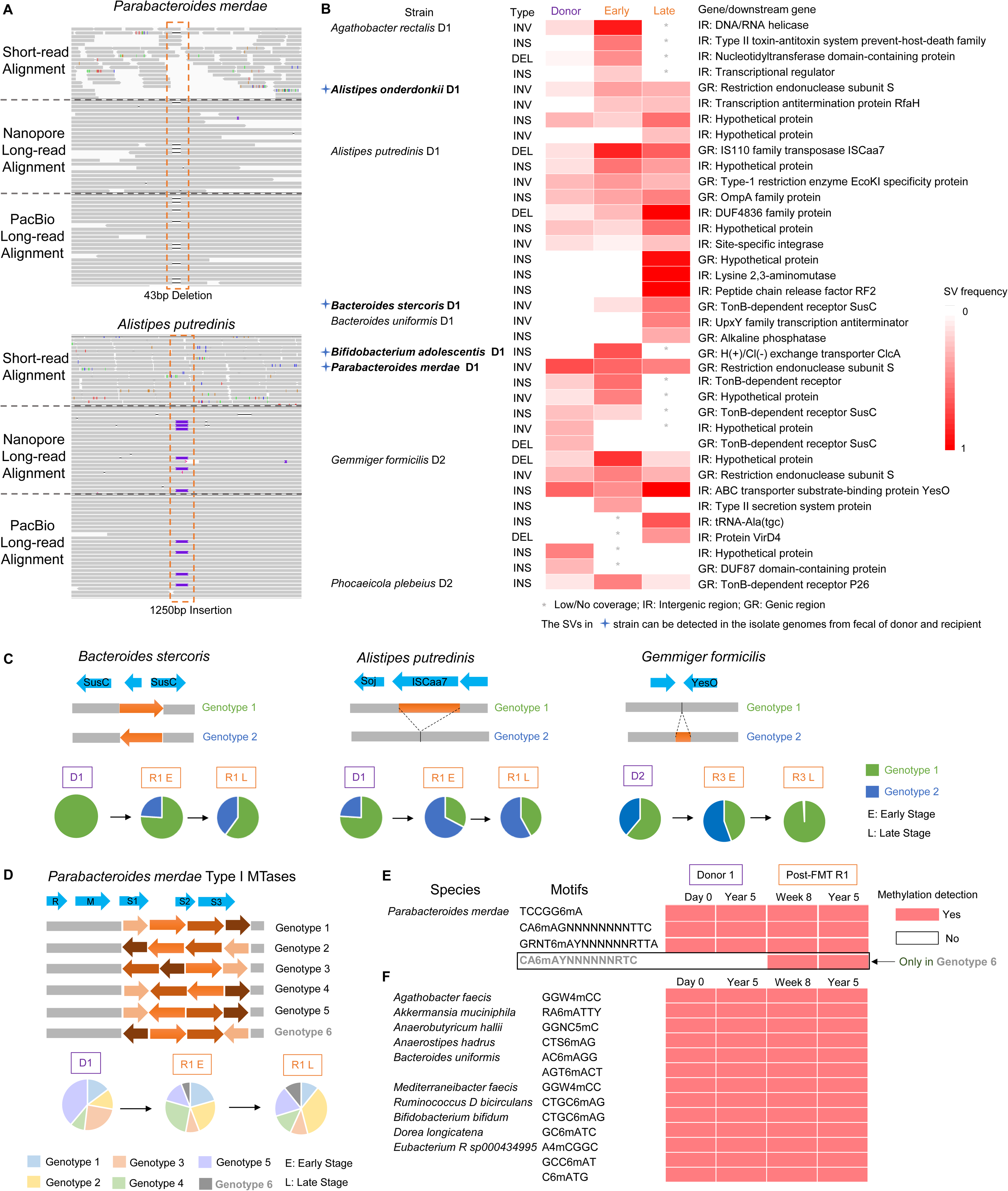
Long reads uncovered complex structural variations in engrafted strains during the 5-year follow-up after rCDI FMT. **A**, IGV plots illustrating the advantage of long reads in the detection of intra-strain structural variation (SV) detection, including a 43 bp deletion in *Parabacteroides merdae* and a 1250 bp insertion in *Alistipes putredinis*, over short-read sequencing. Flanking sequences of SVs show clear and consistent alignments in all long-read IGV plots, supporting that they are from the same strain. In contrast, short-read plots display several mismatches, making it difficult to reliably determine whether the sequences belong to the same strain. **B**, Heatmap summarizing SVs detected in donors and post-FMT recipients at early (E) and late (L) time points after FMT including inversions (INV), deletions (DEL), and insertions (INS). Samples with insufficient coverage over a specific SV region are marked by an asterisk (*). SVs confirmed in isolates from both donor and recipient are marked with a blue cross (+). SVs are also categorized by their location: intergenic regions (IR) and genic regions (GR). **C**, Examples of SVs with changing genotype frequencies across donor and post-FMT recipient over time. (1) inversion in *Bacteroides stercoris*; (2) deletion in *Alistipes putredinis* changes in genotype frequency both in the early and late stage post-FMT; (3) insertion in *Gemmiger formicilis* with frequency changes only at late stage post-FMT. Pie charts demonstrate the genotype frequencies for donor (D1 or D2) and post-FMT recipient (R1 or R3) at early and late time points after FMT. **D**, A complex inversion (shufflon) involving six genotypes is located on restriction endonuclease subunit S of the RMS system in *Parabacteroides merdae*. Genotype frequencies vary with major changes observed for Genotype 6 *i.e.* absence in donor and increases after FMT. **E**, DNA adenine methylation (6mA) motifs in the *P. merdae* strain detected across donor and post-FMT recipient with different time points; CA6mAYNNNNNNRTC corresponds to Genotype 6 of this RMS phase variant, which is only detectable after FMT. **F**, DNA methylation motifs detected across several single strains demonstrating epigenomic stability in the donor (D1) and post-FMT recipient (R1) over time.

Notably, five SVs were identified in TonB-dependent receptor genes or their upstream regions (**Fig. 5B**). which are crucial for iron transport in Gram-negative^73^. The deletions in these genes might be associated with immune-mediated selective pressures^74^. *Bacteroides stercoris* D1 exhibited a high-frequency inversion exclusively in susC genes in late-stage post-FMT, potentially linked to immune adaptation within new host environment (**Fig. 5C**). *A. putredinis* D1, a deletion in IS110 family transposase gene (ISCaa7) was observed with a high prevalence in post-FMT (**Fig. 5C**), and may affect genetic diversity, evolution, and antibiotic resistance propagation, highlighting the impact on microbial adaptability^75,76^. Moreover, in *Gemmiger formicilis* D2, an insertion proximal to yesO gene (**Fig. 5C**), encoding an ABC transporter substrate-binding protein, which likely plays a significant role in nutrient uptake and metabolic processes^77^.

We also observed some more complex SVs that involve bacterial restriction-modification systems (RMS), which safeguard bacterial genomes by cleaving unmethylated foreign DNA while methylating self DNA^45,78–82^. These SVs can alter the activity or specificity of DNA methyltransferases (MTases), driving genome-wide methylation changes that regulate gene expression and enhance phenotypic diversity^83–85^, promoting bacterial adaptation to stresses such as antibiotics and host immune responses^86,87^. Long-read sequencing facilitates simultaneous detection of SVs and DNA methylation patterns at single-molecule resolution, enabling linkage of RMS genotypes to methylation motifs, revealing the epigenomic impact of SVs. For example, we detected a complex SV spanning a Type I RMS S subunits of *P. merdae.* The nested inverted repeats create a complex genomic shuffling among six different genotypes (**Fig. 5D**). Notably, genotype 5, predominantly observed in the donor, had a drastic reduction in post-FMT, while genotype 6 was absent in the donor but significantly increased in post-FMT. Because the specificity unit, S, of the Type I RMS system determines the sequence recognition motif of the MTases, the genotype switches of this RMS locus can create genome-wide methylation changes^41^. To test this hypothesis, we used PacBio sequencing data, which offers advantages in detecting 6-methyladenine (6mA) at single molecule resolution^46,72,80^, and indeed observed a DNA methylation motif CA6mAYNNNNNNRTC exclusively in post-FMT samples (**Fig. 5E**). This observation suggests some SVs can influence genome-wide changes in methylation and potentially help *P. merdae* adapt to the recipient’s gut environment. In addition, we examined 11 unique Type II methylation motifs across ten single strains with high-coverage Nanopore long-read data (**Fig. 5F**), including one 5-methylcytosine (5mC) motif, two as 4-methylcytosine (4mC) motifs, and eight 6mA motifs. These motifs demonstrate consistency and stability in both donors and recipients over various time points.

## Discussion

In this study, we present a novel approach, LongTrack, that uses long-read sequencing and tailored informatics for precision FMT strain tracking, addressing a critical gap in the high-resolution tracking of microbial strains.

A key methodological innovation of LongTrack is the rigorous selection of strain-unique k-mers from long-read MAGs that are compatible with the high-purity yet sometime incomplete long-read MAGs (for low-abundance strains), which are different from separately assembled bacterial isolate genomes^20^. LongTrack includes a critical step that retains only those unique k-mers originating from assembled genomic regions shared across the conspecific strains both within and between FMT donor and recipient samples. Once reliable strain-unique k-mers are selected by LongTrack from long-read MAGs, subsequent strain tracking can then be reliably performed using short-read microbiome sequencing data. This LongTrack framework was validated against ground truth methods based on Strainer with isolate genomes from large-scale bacterial culture^20^.

The previous isolate-based method served as valuable controls for methods evaluation, although standard culture-based approaches often miss certain bacterial taxa due to the selective nature of culture media. For instance, strains like *A. muciniphila* and *F. prausnitzii* were missed under standard culture conditions, despite being culturable from stool samples under more stringent conditions^88,89^. Nonetheless, the inclusion of isolate genomes in our study was primarily intended to evaluate LongTrack with long-read MAGs against Strainer with short-read MAGs, particularly in resolving co-existing strains. Importantly, the use of isolate strains with pure and complete genome assemblies provides a solid approach to evaluate strain tracking^20^. A noteworthy observation from our analysis is that several MAGs with low completeness (<50%) still exhibit high ANI (>99.6%) and k-mer similarity (>92%) to their corresponding cultured isolates. These MAGs, though incomplete, such as *Bacteroides caccae* with MAG size of 499 kb, which is well above the typical size of mobile genetic elements (1–200 kb), contained sufficient unique k-mers to enable accurate strain tracking in post-FMT samples (**Fig. 3C**), and did not impact on the reliability of LongTrack’s results. Application of LongTrack to FMT cases for both rCDI and IBD patients demonstrated the method’s applicability across diverse microbiome complexities. Notably, the application of LongTrack allowed the identification of a large number of strains across diverse taxonomy, including many strains not cultured by standard conditions^20^, and revealed more complete insights into their long-term engraftment in rCDI patients up to five years post-FMT. In IBD FMT, LongTrack demonstrated its effectiveness in more complex microbiomes with greater strain diversity, and the FMT from multiple donors to IBD patients. Due to the relatively small sample size, the study of rCDI and IBD FMT samples were mostly for methods evaluation and demonstration of broad applicability, instead of generalized findings on strain engraftment. Future studies on large cohorts could further explore the influence of strain richness on engraftment dynamics and its therapeutic implications^68^.

The required sequencing depth for effective strain tracking using LongTrack appears to be flexible and influenced by microbial complexity and strain abundance. For donor D1, we generated 23 Gb of long-read data to maximize recovery of strains shared with cultured isolates and enable a robust comparison. However, donors D2 and D3 yielded consistent and accurate reliable tracking results with only 18 Gb and 15 Gb of data, respectively. Notably, in the two IBD FMT cases (D4 and D5), sequencing depths of just 10 Gb and 6 Gb enabled the successful reliable tracking of 52 and 39 strains. These results highlight that moderate sequencing efforts can still yield highly informative outcomes.

Beyond identifying strains that are stably engrafted post-FMT, this study made the first attempt to assess the genomic dynamics of donor strains as they adapt to new microbial and host environments in recipients. This adaptation process, which may select for certain genomic changes in the engrafted strains, remained a relatively unexplored area in FMT research. A number of recent studies highlighted the importance of SVs, in addition to SNVs, in microbiome adaptation^66,81,82^. However, reliable detection of strain-level SVs, across the longitudinal samples poses a challenge to short-read sequencing but can be better approached by long-read sequencing^83^. Long reads have the advantages in detecting complex SVs like shufflons, dynamic DNA inversion systems known to influence bacterial adaptation by generating structural diversity^81,92,93^. Building on longitudinal samples collected over five years following FMT, we identified 38 SVs using stringent filtering criteria (requiring >5 coverage depth for high-confidence mapping) and note that the SV analysis is dependent on sequencing depth and additional SVs may be detected with deeper sequencing data.

We employed two long-read sequencing technologies: Oxford Nanopore and PacBio sequencing. Oxford Nanopore technology produces ultra-long reads that are highly effective for *de novo* assembling high-quality MAGs and detecting SVs. PacBio sequencing offers the advantage of high-accuracy long read and quantitative detection of 6mA and 4mC at single-molecule resolution^72^. By leveraging the complementary strengths of these technologies, we suggest a new dimension to characterize how donor strains adapt in post-FMT recipients over time.

While LongTrack offers novel insights, it does inherently face limitations in recovering super low abundance strains. To mitigate this, we propose two complementary strategies. First, integrating long-read sequencing and culture-based methods to provide a more complete view of the FMT strain engraftment. Second, applying targeted enrichment methods, such as flow cytometry^84^ or bacterial methylome patterns^85^, to selectively sequence and analyze low-abundance taxa, thereby improving their recovery and tracking.

As the cost of long-read sequencing platforms continue to decrease, we expect LongTrack to be broadly used for high-resolution strain tracking in FMT studies, paving the way towards better understanding FMT and the development of more effective defined biotherapeutics. More broadly, LongTrack can also help precise strain tracking in other biomedical studies to enhance our understanding of strain transmission, genomic and epigenomic variation, and their roles in health and diseases^95–98^.

## Extended data figure legend

**Extended Data Fig. 1 Description of sample collection and FMT study design for CDI patients.**

Stool samples were collected from three donors and four recipients at multiple time points. For the illustration of LongTrack, strain tracking was performed for long-read MAGs assembled from donors and recipients using short-read metagenomic data generated across all the samples shown in the figure (please refer to **Methods** for details). The Venn diagram displays the number of strains that were recovered by long-read (LR) MAGs (yellow color) or isolate genomes from bacterial culture (red color) from 3 donors and 2 recipients

**Extended Data Fig. 2 Illustration of LongTrack workflow’s implementation in this study for generating strain-level long-read MAGs**

Initially, the raw long reads from donor and pre-FMT recipient underwent metagenomic assembly using the metaFlye (v2.9). The resulting Metagenome-assembled genomes (MAGs) were then partitioned into individual bins corresponding to the species level using MaxBin (v2.0). Strainberry (v1.1) was used to separate the contigs into strain level based on haplotype phasing for each species-level bin. Subsequently, metaWRAP (v1.3.2) was employed to bin the separated contigs into strain level. Finally, the strain-level long-read MAGs with completeness of more than 10% were chosen for FMT strain tracking in post-FMT recipients (Please see **Methods** for details).

**Extended Data Fig. 3 Comparative evaluation of long-read MAGs and short-read MAGs from Donor 1**

**A.** Completeness levels of 124 long-read MAGs (LR MAGs) and 135 short-read MAGs (SR MAGs) from Donor 1 reveal no significant differences. **B**. Contamination levels of 124 LR MAGs and 135 SR MAGs from Donor 1 assessed using CheckM, showing significantly lower contamination in LR MAGs (p < 0.01, **, Wilcoxon test). **C**. Contig N50 (kb) comparison for 124 LR MAGs and 135 SR MAGs Donor 1.

**Extended Data Fig. 4 Assessment of contamination level in long-read/short-read MAGs**

**A.** Schematic of contamination level by mapping long-read and short-read MAG to its corresponding isolate genome, calculated as the percentage of unaligned bases contamination; **B.** Comparative analysis of contamination levels determined by comparing to the isolate genome and by using the CheckM tool. A total of 60 long-read (LR) MAGs from Donors 1–3 and Recipients 1–4, and 23 short-read (SR) MAGs from Donor 1 (D1) were analyzed. (**C–D**) Comparison of contamination levels in corresponding strains between LR MAGs and SR MAGs from D1. (**C**) Bar plots illustrating contamination levels in corresponding strains between LR MAGs and SR MAGs for species with multiple co-existing strains and species with a single strain, emphasizing the consistently lower contamination observed in LR MAGs. **D.** Contamination levels for 31 LR MAGs and 23 SR MAGs from D1, categorized by species with either a single strain or multiple co-existing strains. LR MAGs consistently demonstrated lower contamination levels (p < 0.01, permutation test). **E.** Genomic similarity of shared strains between the long-read (LR) MAGs/short-read (SR) MAGs and their corresponding isolate genomes determined by k-mer similarity (**Methods**). The result indicates long-read MAGs exhibit a significantly higher similarity (*p-value* < 0.05, *).

**Extended Data Fig. 5 Evaluation of FMT strain tracking between LongTrack and StrainFinder as a representative of SNP-based inference approach**

**A.** For the donor sample D1, *in silico* spike-in dataset was designed with 3% *Clostridium tyrobutyricum* and 2% *Turicibacter sanguinis* added to the microbiome dataset. For the post-FMT recipient R1, the spike-in data set A (low abundance) had a composition of 0.25% *Clostridium tyrobutyricum* and 0.1% *Turicibacter sanguinis*, while the spike-in data set B (high abundance) had a composition of 2.5% *Clostridium tyrobutyricum* and 1% *Turicibacter sanguinis* **(Methods)**. **B**. The strain tracking results indicate that the SNP-based approach, StrainFinder, works well for strains with relatively modest-to-high abundance (data set B), but tends to miss strains with modest-to-low abundance (data set A). The chart represents the presence (green) or absence (gray) of strains in post-FMT recipients based on spike-in design (considering ground truth), LongTrack with long-read MAGs, or StrainFinder. Discrepancies with the ground truth are indicated by an "X" marker.

**Extended Data Fig. 6 Sensitivity evaluation of LongTrack and culture-based approach (Strainer)**

The sensitivity evaluation of LongTrack and the culture-based approach (Strainer) was performed using the union of all strains detected either through isolate genomes or MAGs as the reference set. To evaluate sensitivity across different microbial abundance levels, the reference set was stratified by relative abundance thresholds, which provides a comprehensive benchmark that encompasses the diversity of detectable strains, including those uniquely identified by each method.

**Extended Data Fig. 7 LongTrack uncovered engraftment of cultured strains across FMT cases in rCDI patients**

**A**, Proportional engraftment of donor strains (PEDS) for shared cultured strains is shown across individual post-FMT recipients at various time points of the 5-year follow up. **B**, Proportional persistence of recipient strains (PPRS) for shared cultured strains is shown across individual post-FMT recipients at various time points of the 5-year follow up.

**Extended Data Fig. 8 LongTrack uncovered hundreds of engrafted strains of rCDI**

The long-read MAGs uncultured by the previous study were utilized for strain tracking to determine if they were present in the post-FMT recipients for FMT cases of CDI: The results of tracking across three FMT cases D1->R1 (**A**), D2->R3 (**B**), and D3->R4 (**C**), which describe the presence (green) or absence (gray) of strains based on LongTrack.

**Extended Data Fig 9. Reverse strain tracking of earlier samples based on long-read MAGs assembled from post-FMT recipients using LongTrack**

The long-read MAGs assembled from post-FMT recipient samples were utilized for strain tracking to determine if they were also present in the donor and pre-FMT recipient metagenomic data using LongTrack (Reverse tracking). The results of reverse tracking across three FMT cases: D1 -> R1, D2 -> R3 and D3 -> R4 are shown in **A-C**, which describe the presence (green) or absence (gray) of strains based on LongTrack. **(D-F)** Illustration of bacterial strains that may have been acquired from environmental sources, as determined by long-read MAGs assembled from the five-year post-FMT recipients’ samples but absent in all the earlier samples. The results across three FMT cases: D1 -> R1, D2 -> R3 and D3 -> R4 are shown in **D-F**, which describe the presence (green) or absence (gray) of strains based on LongTrack.

**Extended Data Fig 10. Description of samples collection and FMT study design for two IBD patients.**

Stool samples were collected from six donors and two recipients at multiple time points. The two recipients (IBD R5 and IBD R6; orange) had received FMT from six donors (IBD D4 - D9; gray). For the illustration of LongTrack, strain tracking was performed for long-read MAGs assembled from two donors (IBD D4 and D5; purple) using short-read metagenomic data generated across all the samples shown in the figure (please refer to **Methods** for details)

## Acknowledgments

This work was supported in part by the staff and resources of the Microbiome Translational Center and Department of Scientific Computing at the Icahn School of Medicine at Mount Sinai. This work was supported by grant no. R35 GM139655 (G.F.) from the National Institutes of Health. G.F. is a Hirschl Research Scholar by Irma T. Hirschl/Monique Weill-Caulier Trust and a Nash Family Research Scholar.

## Competing interests

S.P. has served as a consultant / steering committee member for Vedanta Biosciences and has received speaker / advisory board fees from AbbVie, Dr Falk Pharma, Ferring, Janssen and Takeda. J.F. is a consultant for Vedanta Biosciences and Genfit. The remaining authors declare no competing interests.

## Author contributions

Initiated study: G.F. and J.F. Developed methodology: Y.F. and G.F. Data analysis: Y.F., J.F. and G.F. Data interpretation: Y.F., M.N., V.A., E.A.M., M.K., L.C., J.F. and G.F. Collection and interpretation of clinical data: M.A.K., T.J.B., S.P., N.O.K. and A.G. Supervised research: G.F. Wrote first draft of paper: Y.F. and G.F. Approved paper: all authors.

## Data availability

All sequencing data generated for this study have been uploaded to the Sequence Read Archive under the BioProject PRJNA1101882.

## Code Availability

Customized scripts of LongTrack and a tutorial with supporting data are available at https://github.com/fanglab/LongTrack

## Materials and Methods

### Fecal sample collection and DNA extraction

The adult fecal samples were from a previous study^1^ with Institutional Review Board approval (rCDI: HS# 11-01669, IBD: HREC/13/SVH/69). Fecal DNA was extracted with a QIAamp PowerFecal Pro DNA Kit (Qiagen, 51804) according to the manufacturer’s instructions. Briefly, 64 - 436 (average is 181) mg of fecal sample, and 800 μl of Solution CD1 were added to the Bead Tube (provided). After gentle vortexing, the sample was homogenized with the TissueLyser II machine (Qiagen, 85300) for 10 min at 25 Hz speed (5 min each side). The homogenized samples were centrifuged to collect the supernatant at 15,000 x g for 1 min. The subsequent column-based extraction steps were performed as described in the QIAamp PowerFecal Pro DNA Kit protocol.

### Library preparation and sequencing

#### Nanopore long-read sequencing

Native libraries were prepared using the Oxford Nanopore Technology Inc. (New York, NY, USA) Native barcoding kit (EXP-NBD104 and 114 for R9.4.1 flow cells, and SQK-NBD114.24 for R10.4.1 flow cells), Ligation Sequencing Kit (SQK-LSK109 for R9, and SQK-LSK114 for R10.4.1). Prior to library preparation, genomic DNA (gDNA) was sheared using Covaris g-tubes (Covaris, Woburn, MA, USA 520079) at 6000 RPM in an Eppendorf 5424R centrifuge (Eppendorf, Enfield, CT, USA, 5406000640) and cleaned up with AMPure XP beads (Beckman Coulter, Indianapolis, IN, USA, A63882) at 0.8 x concentration. Whole genome amplification (WGA) samples were prepared using REPLI-g Mini Kits (QIAGEN, Hilden, Germany, 150023) according to the protocol with 12.5 ng of input DNA per sample and 16 h incubation at 30 °C, followed by denaturation at 65 °C for 3 minutes. Next, WGA samples were treated with T7 endonuclease I (NEB M0302S) to maximize nanopore sequencing yield according to ONT documentation, then cleaned up with AMPure XP beads as above. Subsequently, samples were processed the same as above native libraries. Samples were sequenced on R9.4.1 and R10.4.1 MinION flow cells and post-run basecalled using Guppy (v5.0.7 for R9.4.1 data and 6.3.4 for R10.4.1 data)^2^ with the SUP model dna_r9.4.1_450bps_sup.cfg and dna_r10.4.1_e8.2_400bps_sup.cfg, respectively.

#### Illumina short-read sequencing

Library construction for Illumina sequencing was executed utilizing the seqWell Plexwell 384 kit. DNA was systematically barcoded, and ligation products underwent purification. Size selection and PCR amplification of the DNA fragments followed. The prepared library was then sequenced on an Illumina HiSeq platform, generating paired-end reads of 150 base pairs (bp) each.

#### PacBio long-read sequencing

DNA SMRTbell libraries were prepared according to the manufacturer’s instructions for the SMRTbell Express Template Prep Kit 2.0, then each library was annealed to sequencing primer together with the Sequel 2.1 polymerase, followed by loading to the a 8M Sequel II sequencing chip (for Sequel II) and 25M Revio sequencing chip.

### Metagenomic assembly, strain-level separation and taxonomic annotation

#### Long-read metagenomic assembly

Initially, the raw long reads were subjected to a metagenomic assembly for long-read MAGs process using the metaFlye tool (v2.9)^3^ with “---nano-raw --meta --keep-haplotypes --iterations 2”. The resulting metagenome-assembled genomes (MAGs) were partitioned into individual bins corresponding to the species level using the MaxBin (v2.0) software^4^. To annotate the species-level bins, we applied the Genome Taxonomy Database Toolkit (GTDB-tk v1.6.0)^5^. For each species-level bin, we utilized Strainberry (v1.1)^6^ to separate the contigs into the strain level based on haplotype phasing. Subsequently, these separated contigs were binned to strain level using the "binning" module of metaWRAP (v1.3.2)^7^, which could combine the results from three metagenomic binning software - MaxBin2^4^, metaBAT2^8^, and CONCOCT^9^. The quality of the reconstructed long-read MAGs was evaluated using the "bin_refinement" module in metaWRAP (v1.3.2)^7^, which combines the results of multiple binning algorithms and selects high-quality bins based on completeness and contamination estimates, as well as estimated by CheckM (v.1.1.3)^10^. The strain-level long-read MAGs with completeness of more than 10% were selected for further analysis. We used viralVerify (v1.1)^11^ and PLASMe (v1.1)^12^ to identify plasmids in each assembly. For viralVerify, only contigs shorter than 500 kbp with prediction scores greater than five were considered. Annotations labeled as ‘Plasmid’ or ‘Uncertain - plasmid or chromosomal’ were classified as plasmids. For PLASMe, only results with high precision were included. Overlapping plasmids identified by both methods were treated as the final results.

#### Short-read metagenomic assembly

We conducted a metagenomic assembly of the short reads using metaSPAdes (v3.15.2)^13^ and MEGAHIT (v1.1.3)^14^ with default settings to construct short-read MAGs, which show similar completeness and contamination levels (**Supp Figs. S1A and S1B).** However, MEGAHIT^14^ identified a smaller number of shared strains with isolate genomes (**Fig 2B**, **Supp Fig. S1C).** Based on these results, we used MAGs from metaSPAdes^13^ for subsequent analyses.

Subsequently, we utilized metaWRAP (v1.3.2)^7^ to bin and refine the reconstructed short-read MAGs. The completeness and contamination level for the short-read MAGs were estimated using CheckM (v.1.1.3)^10^, and only those with completeness of more than 10% were selected for further analysis. Taxonomic assignment of the reconstructed short-read MAGs was performed using GTDB-tk (v1.6.0)^5^.

### Evaluation of long-read MAGs

The previous study identified a total of 184 unique strains, which were from 184 different bacterial isolates previously isolated from three distinct donors and recipients^1^. Specifically, 136 bacterial isolates were from Donor 1 (D1), 20 were from Donor 2 (D2), 8 were from Donor 3 (D3), 2 were from Recipient 1 pre (R1), 7 were from Recipient 4 pre (R3), and 11 were from Recipient 4 pre (R4). The resulting assemblies for each isolate serve as reference individual genomes. To detect shared strains between long-read MAGs/short-read MAGs and genomes of bacterial isolates, we utilized fastANI (1.32)^15^ for average nucleotide identity (ANI) calculation, applying a 99.4 % cut-off^3^. We assessed the contamination level by mapping long-read and short-read MAG to its corresponding isolate genome, calculated as the percentage of unaligned bases. Mummer (v4.0.0b)^17^ was employed to identify the mapping regions between long-read MAG/short-read MAG and its corresponding genome of isolate. Additionally, permutation tests were performed to evaluate statistical differences in contamination levels between long-read and short-read MAGs. We also assessed the genomic similarity of shared strains between the long-read MAGs/short-read MAGs and their corresponding isolate genomes determined by k-mer similarity as in the previous study^1^, with the following formula:

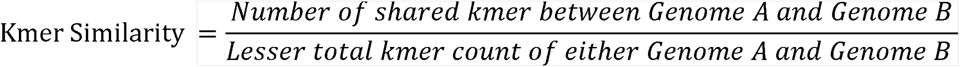

Where:

K-mer size is set to 31.
Shared k-mers are those that appear in both Genome A and B.
Total k-mer count is the count of all distinct 31-mers in a genome.

The k-mer similarity between two genomes A and B was determined by generating a hash for genomes with k-mer size 31 and quantifying the proportion of k-mers shared in both genomes A and B divided by the lesser total k-mer count of either

### The proportion of reads associated with each strain/MAG

We analyzed long-read and short-read metagenomic sequencing data from D1 by mapping to a merged set of genomes, which included genomes from bacterial isolates and uncultured long-read MAGs or short-read MAGs. For the long-read data, we used 136 genomes from bacterial isolates and 105 uncultured long-read MAGs. The proportion of reads associated with each strain/MAG was determined by calculating the percentage of metagenomic reads that were successfully mapped to each genome/MAG in this merged set. Similarly, the short-read data were aligned to the same 136 genomes of bacterial isolates, supplemented with 120 uncultured short-read MAGs, to quantify the proportion of reads associated with each strain/MAG.

### LongTrack framework designation for FMT strain tracking based on strain-unique k-mers

The LongTrack framework for FMT strain tracking follows a three-step algorithmic process (**Fig. 1C**). LongTrack strain tracking relies on strain-unique k-mers, built on previous methods^1^, and incorporates important methodological adaptations compared to previous studies to ensure the reliable selection of unique k-mers and to minimize false positives.

#### Step 1: Identification of strain-unique k-mers from each long-read MAG

##### a. Extract strain-unique k-mers from each long-read MAG

Following established methods^1^, the LongTrack framework extracts k-mers (k=31) for each strain-level long-read MAG from the donor or pre-FMT recipient using KMC (version 3.0.0)^18^. Strain-unique k-mers are then identified by eliminating common k-mers shared in 100,000 bacterial genomes in the NCBI database (as of 2018-11-20), 116 unrelated metagenomic samples from a previous study^1^, and other long-read MAGs within the same FMT case.

##### b. Obtain strain-unique k-mers in co-existing strains

Relying solely on strain-unique k-mers identified in Step 1a can lead to false strain-tracking results due to the varying completeness of long-read MAGs, especially for similar co-existing strains. For instance, as shown in **Fig. 1C**, Strains A1 and A2 both belong to Species A and are co-existing strains from the same donor. If the k-mers identified for Strain A1 in Step 1a correspond to genomic regions missing in Strain A2 due to incomplete assembly, these k-mers may also appear in Strain A2 simply because of the high genomic similarity between the two strains. In such cases, the k-mers are not truly unique to Strain A1 but are instead masked in Strain A2 by assembly gaps, leading to unreliable results.

To address this issue, LongTrack introduces methodological adaptations to ensure the reliable selection of strain-unique k-mers and to minimize false positives. Specifically, for similar co-existing strains within a single fecal sample or closely related strains found separately in the donor and pre-FMT recipient samples, LongTrack applies a filter to retain only those unique k-mers originating from genomic regions shared among multiple co-existing strains in same FMT case. Mummer (v4.0.0b)^17^ is used to identify these common genomic regions.

This approach prevents the erroneous selection of k-mers that might seem strain-specific but are unreliable due to MAG incompleteness—a critical distinction between high-purity long-read MAGs and complete genomes from bacterial isolates. The resulting retained strain-unique k-mers serve as markers for each long-read MAG, enabling reliable strain tracking in subsequent steps.

#### Step 2: Assign short reads with final strain-unique k-mers to metagenomic samples

In the second step, the final strain-unique k-mers identified in Step 1b were utilized to assign short reads from post-FMT recipient microbiome data to their corresponding long-read MAGs. Short reads containing strain-unique k-mers were mapped back to their respective long-read MAGs using Bowtie 2.4.1^19^, with parameters optimized for accuracy (‘very-sensitive,’ ‘no-mixed,’ and ‘no-discordant’). The successful alignment of short reads to genomic regions containing strain-unique k-mers confirmed the presence of the corresponding MAG in the post-FMT recipient, facilitating accurate strain tracking.

#### Step 3: Discard results with low confidence score

To ensure the reliability of strain tracking, we calculated a confidence score by comparing read enrichment in five unrelated metagenomic samples (negative controls) as done in the previous study^1^. Results with low confidence scores (< 95% confidence) were discarded to maintain the accuracy of strain tracking.

### Evaluation of LongTrack for FMT strain tracking

#### Sensitivity evaluation of LongTrack and culture-based approach (Strainer)

To evaluate the sensitivity of LongTrack and the culture-based approach (Strainer), we used the union of all strains detected either through isolate genomes or MAGs as the reference set. This method offers a comprehensive benchmark that captures the diversity of detectable strains, including those uniquely identified by each technique. The analysis was stratified by relative abundance thresholds to assess detection performance across varying microbial abundance levels, yielding valuable insights into the sensitivity of LongTrack. Sensitivity was determined as the ratio of True Positives (TP) to the sum of True Positives (TP) and False Negatives (FN).

#### Evaluation of LongTrack for FMT strain tracking through comparison with ground truth

For a systematic assessment of LongTrack for FMT strain tracking, we utilized the shared strains between long-read MAGs/short-read MAGs and genomes of bacterial isolates in four FMT cases of rCDI. This evaluation was conducted by comparing tracking results in the post-FMT recipients at different time points based on Strainer with genomes of bacterial isolates (considered the ground truth) and LongTrack/Strainer with short-read MAGs. We applied two key metrics:

1. Precision: Precision quantifies the proportion of correct tracking results (True Positives, TP) relative to all tracking results predicted by the LongTrack/Strainer with short-read MAGs. It is calculated as TP divided by the sum of True Positives (TP) and False Positives (FP).
2. Specificity: Specificity evaluates the effectiveness of LongTrack/Strainer with short-read MAGs in accurate strain tracking. It is determined by the ratio of True Negative (TN) to the total number of True Negative (TN) and False Positives (FP).

The accuracy of strain tracking based on LongTrack/Strainer with short-read MAGs was evaluated using the formula: the number of consistent tracking results between ground truth and LongTrack/Strainer with short-read MAGs divided by the total number of tracking results, expressed as a percentage. The tracking results were presented as a heatmap using the R package "pheatmap v1.0.12".

#### Evaluation of FMT strain tracking between LongTrack and StrainFinder, which serves as a representative methods of SNP-based tracking

To evaluate FMT strain tracking between LongTrack and StrainFinder^20^, we utilized *in silico* spike-in datasets (**Extended Data Fig. 5**). For donor D1, we spiked in 3% *Clostridium tyrobutyricum* at 60x depth and 2% *Turicibacter sanguinis* at 40x depth into long-read microbiome dataset. The simulated Nanopore long reads were generated using nanosim (v 3.0.0)^21^ for metagenomic assembly. The error profile was trained using real sequencing data from a large number of nanopore sequencing reads, and the read length distribution was modeled to match that of the actual Donor 1 sequencing data. Similarly, we produced simulated short-read data with identical abundances of these strains for donor D1 in the short-read microbiome datasets. For the post-FMT recipient R1, we spiked in short reads to the short-read metagenomic sequencing data, which was used for strain tracking. We designed two distinct spike-in datasets: A) The first dataset A (low abundance) featured a composition of 0.25% (5x depth) *Clostridium tyrobutyricum*, along with paired short reads, each having a length of 150 bp. Furthermore, this dataset included 0.1% (2x depth) *Turicibacter sanguinis*, with paired short reads, also 150 bp in length. B) The second dataset B (high abundance) presented a composition of 2.5% (50x depth) *Clostridium tyrobutyricum*, accompanied by paired short reads, each measuring 150 bp in length. Additionally, this dataset contained 1% (20x depth) *Turicibacter sanguinis*, once again with paired short reads, each 150 bp in length. We simulated paired short reads with a length of 150 bp using randomreads.sh from the bbmap package (v38.86)^22^. Subsequently, LongTrack and StrainFinder^20^ were performed for strain tracking analysis.

#### Evaluation of LongTrack for FMT strain tracking when there are high-similarity strains present in between the donor and pre-FMT samples

To evaluate LongTrack/Strainer with short-read MAGs for FMT strain tracking when there are high-similarity strains present in between the donor and pre-FMT samples, we spiked-in ten pairs of strains, each with high similarity (ANI>99.5%), from ten different species, one strain to D1 and the other to R1 pre-FMT, similarly for all the ten pairs **(Fig. 3F)**. All twenty strains were added to R1 post-FMT. To facilitate metagenomic assembly, we generated simulated long-read data/short-read data for D1 and R1 pre-FMT into their long-read/short-read microbiome datasets. Furthermore, we simulated short-read paired reads, each with a length of 150 bp, to R1 post-FMT short-read microbiome datasets. The simulated Nanopore long reads were generated using the nanosim (v 3.0.0)^21^. The short reads were generated using the randomreads.sh from the bbmap package (v38.86)^22^. LongTrack/Strainer with short-read MAGs was performed for strain tracking analysis.

#### Evaluation of LongTrack for FMT strain tracking when there are co-existing strains of the same species in one donor sample

To evaluate LongTrack/Strainer with short-read MAGs for FMT strain tracking when there are co-existing strains of the same species in one donor sample, we spiked-in three strains of *Phocaeicola massiliensis* (Strains 1-3) to D1 and a fourth strain of *P. massiliensis* (Strain 4) to R1 pre-FMT **(Fig. 3H)**. For metagenomic assembly, long-read data/short-read data were generated for these strains in D1 and R1 pre-FMT microbiome datasets. Furthermore, we simulated short-read paired reads of 4 strains (Strains 1-4), each with a length of 150 bp, to R1 post-FMT short-read microbiome datasets. The simulated Nanopore long reads with an average length of 5 kb, were generated using the nanosim (v 3.0.0)^21^. The short reads were generated using the randomreads.sh from the bbmap package (v38.86)^22^. LongTrack/Strainer with short-read MAGs was performed for strain tracking analysis.

### Application of LongTrack: strain tracking in rCDI and IBD cases, reverse tracking and identification of novel strains

#### Application of LongTrack in strain tracking of rCDI cases

We initially retrieved long-read MAGs from 3 donors and 4 pre-FMT recipients. Using LongTrack, we examined whether these MAGs were present in the metagenomic sequencing data of the corresponding post-FMT recipients across multiple time points.

#### Application of LongTrack in strain tracking of IBD cases

In the IBD cases, the recipients (R5 and R6) had received fecal microbiota transplantation from six donors (D4 - D9), and strain tracking was performed for long-read MAGs from two donors (D4 and D5). To avoid the influence of common k-mers from the remaining four donors, we further excluded the k-mers that were present in the short-read metagenomic sequencing data of the other four donors. Subsequently, we assessed whether these MAGs were present in the metagenomic sequencing data of the corresponding post-FMT recipients across multiple time points using LongTrack.

#### LongTrack-based reverse tracking and novel strain identification

We initially retrieved 402 uncultured long-read MAGs from post-FMT recipients (R1-R4). We then identified long-read MAGs shared between the post-FMT recipient and the respective donor or pre-FMT recipient using an ANI exceeding 99.4% in the same FMT cases. We confirmed that 39 long-read MAGs are unique to the post-FMT recipients and investigated whether they were present in the metagenomic sequencing data of the corresponding donor and pre-FMT recipient using LongTrack. If the long-read MAGs were not tracked in microbiome data of donor, pre-FMT or any earlier time points of post-FMT, we considered them to be novel strains.

### Detection of structural variation in engrafted strains using long-read sequencing

We utilized NGMLR (v 0.2.7)^23^ to perform the alignment of long-read microbiome data of post-FMT recipients (R1-R4) across various time points to the uncultured long-read MAGs and isolated genomes from the respective rCDI donors (D1-D3). Following the alignment process, we employed Sniffles (v1.0.12)^24^ for the detection of structural variations (SVs) within the long-read microbiome data. To ensure the reliability of our SV detection, we applied several filtering criteria: 1) We excluded SVs involving multiple strains within the same species, as these are more prone to mismatches and may not accurately represent structural variations. 2) For mapping quality control, we retained aligned reads with identity greater than 98%, length more than 1000 bp, and MAPQ (Mapping Quality) score greater than 60. 3) We kept SVs that had a coverage depth of greater than 5 in long-read metagenomic sequencing data of donor and at least one post-FMT, ensuring that the detected structural variations were well-supported by sequencing data.

### Validation of structural variations using isolate long-read sequencing

We collected 18 isolates for SV validation, including:

- 2 *Alistipes onderdonkii* isolates: 1 from the donor and 1 from 8 weeks post-FMT.
- 4 *Bacteroides stercoris* isolates: 2 from the donor, 1 from 8 weeks post-FMT, and 1 from 6 months post-FMT.
- 5 *Bifidobacterium adolescentis* isolates: 2 from the donor, 1 from 8 weeks post-FMT, and 2 from 6 months post-FMT.
- 7 *Parabacteroides merdae* isolates: 3 from the donor, 2 from 8 weeks post-FMT, and 2 from 6 months post-FMT.

Native libraries were prepared using the Oxford Nanopore Technologies Native Barcoding Kit (SQK-NBD114.24) for R10.4.1 flow cells and the Ligation Sequencing Kit (SQK-LSK114).

Samples were sequenced on R10.4.1 MinION flow cells, and basecalling was performed post-run using Guppy (v6.3.4)^2^ with the SUP model dna_r10.4.1_e8.2_400bps_sup.cfg.

For alignment, we utilized NGMLR (v0.2.7)^23^ to map the single isolate long-read sequencing data to the corresponding long-read MAGs. SVs were detected using Sniffles (v1.0.12)^24^. We then cross-validated the detected SVs from metagenomic sequencing data to determine if they could also be identified in the single isolate long-read sequencing data.

### Methylation analysis using long read sequencing

#### De novo methylation motif discovery using Nanopore long-read sequencing

We utilized Nanodisco (v1.0.2)^25^ to identify *de novo* methylation motifs in long-read MAGs from the D1. First, raw fast5 reads from both native and WGA metagenomic sequencing data of D1 were aligned to the long-read MAGs with “nanodisco preprocess”. The alignments for each long-read MAG were then segregated. Only long-read MAGs exhibiting a coverage above 20 x were retained to following analysis. We then computed current differences (PA) between the native (retaining methylation) and WGA (methylation-free) data for each genomic position using “nanodisco difference”. To conclude, “nanodisco motif and refine” was employed for *de novo* methylation motif discovery based on these current differences. For D1 5-year, R1 post-FMT 8-week, and R1 post-FMT 5-year, the raw fast5 reads from native microbiome data were aligned to the retained long-read MAGs from D1 with “nanodisco preprocess” and separated the alignments into each MAG. Subsequently, we computed the current differences by comparing the native data with the WGA data from D1 for each MAG using “nanodisco difference”.

“Nanodisco refine” was employed to verify the presence of D1 detected methylation motifs in the samples of D1 5-year, R1 post-FMT 8-week, and R1 post-FMT 5-year based on the current differences.

#### De novo methylation motif discovery using PacBio long-read sequencing

For PacBio long-read data of D1 (generated by the Sequel II platform), the raw reads pre-processing was performed with SMRTlink (v8.0). Raw multiplexed files were first de-multiplexed with lima. We mapped the subreads to long-read MAGs using pbalign (v0.4.1) with default parameters and separated the alignments into each MAG. The IPD ratio was calculated with ipdSummary (v2.4.1) from the alignments for each MAG. The *de novo* methylation motifs were detected by using motifMaker (in SMRTlink v8.0 package) with default parameters. For PacBio long-read data of D1 5-year, R1 8-week and R1 5-year (generated by the Revio platform), the raw reads pre-processing was performed with SMRTlink (v13.0), and methylation motif detection was performed by our in-house scripts.

## Notes

### Summary of Updates

We conducted additional analyses to more precisely evaluate sensitivity and addressed the reviewer's minor suggestions.

