## Extended Data Figures for "Long read metagenomics-based precise tracking of bacterial strains and genomic changes after fecal microbiota transplantation"

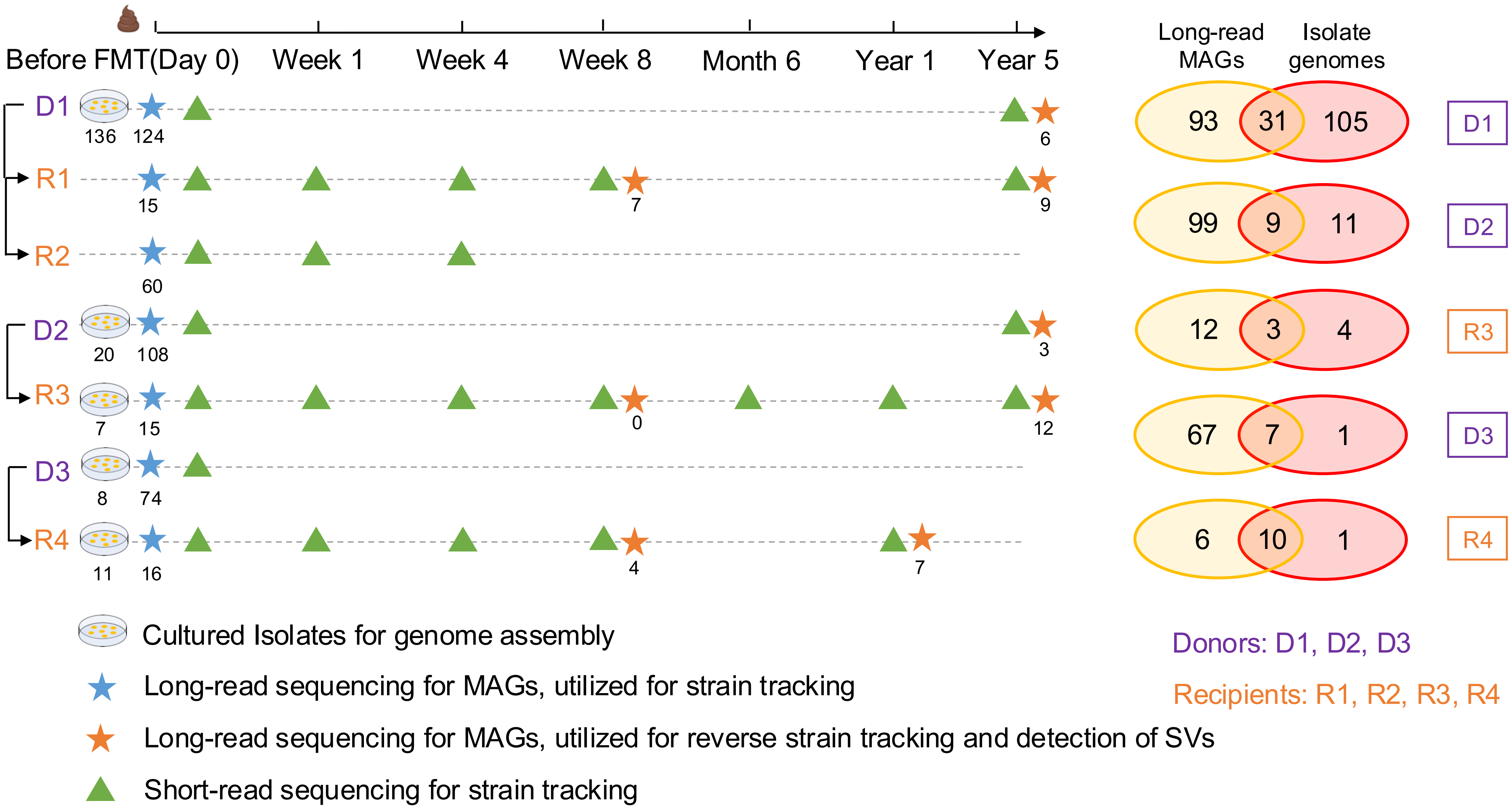
Before FMT(Day 0)
Week 1
Week 4
Week 8
Month 6
Year 1
Year 5
Long-read MAGs
Isolate genomes
D1
136
124
6
R1
15
7
9
R2
60
D2
20
108
3
R3
7
15
0
12
D3
8
74
R4
11
16
4
7
93
31
105
D1
99
9
11
D2
12
3
4
R3
67
7
1
D3
6
10
1
R4
Cultured Isolates for genome assembly
Long-read sequencing for MAGs, utilized for strain tracking
Long-read sequencing for MAGs, utilized for reverse strain tracking and detection of SVs
Short-read sequencing for strain tracking
Donors: D1, D2, D3
Recipients: R1, R2, R3, R4

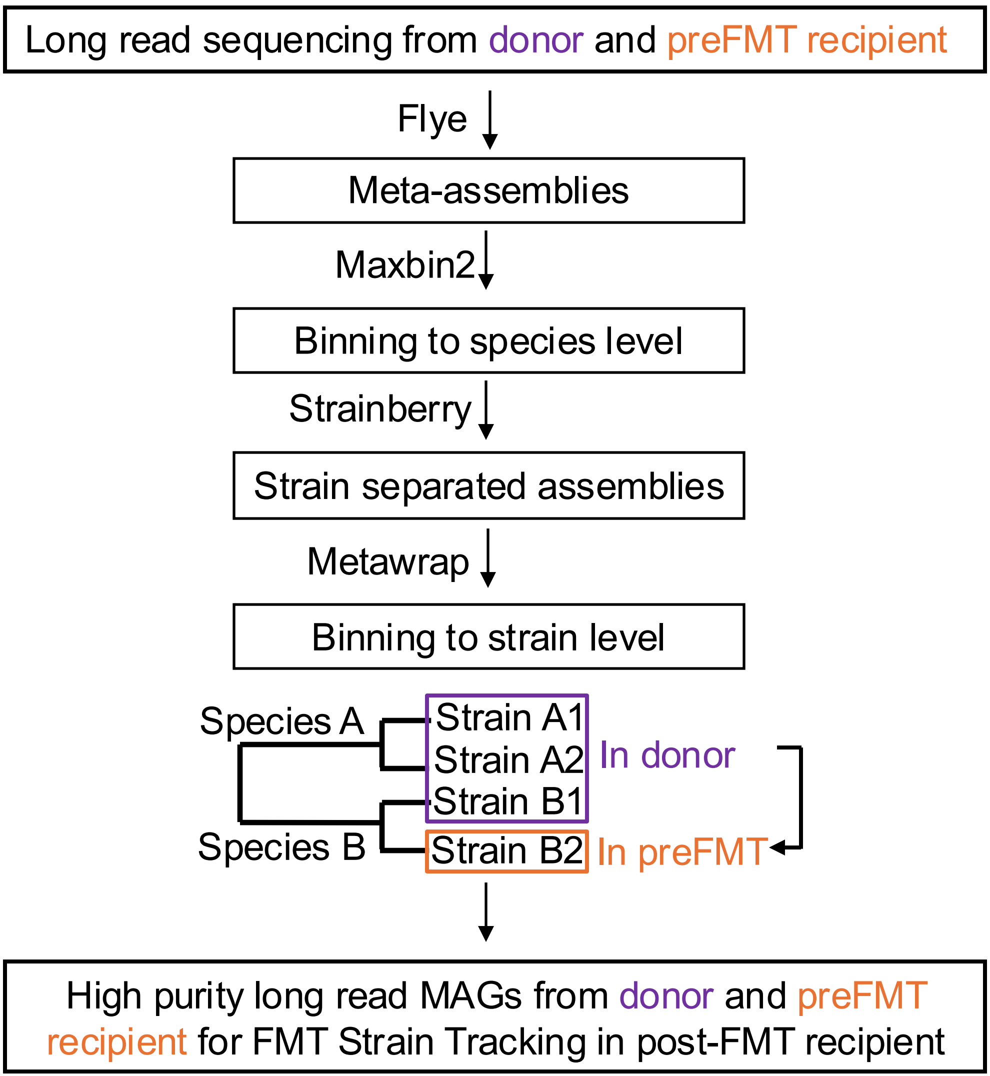
Long read sequencing from donor and preFMT recipient
Flye
Meta-assemblies
Maxbin2
Binning to species level
Strainberry
Strain separated assemblies
Metawrap
Binning to strain level
Species A
Species B
Strain A1
Strain A2
Strain B1
Strain B2
In donor
In preFMT
High purity long read MAGs from donor and preFMT recipient for FMT Strain Tracking in post-FMT recipient

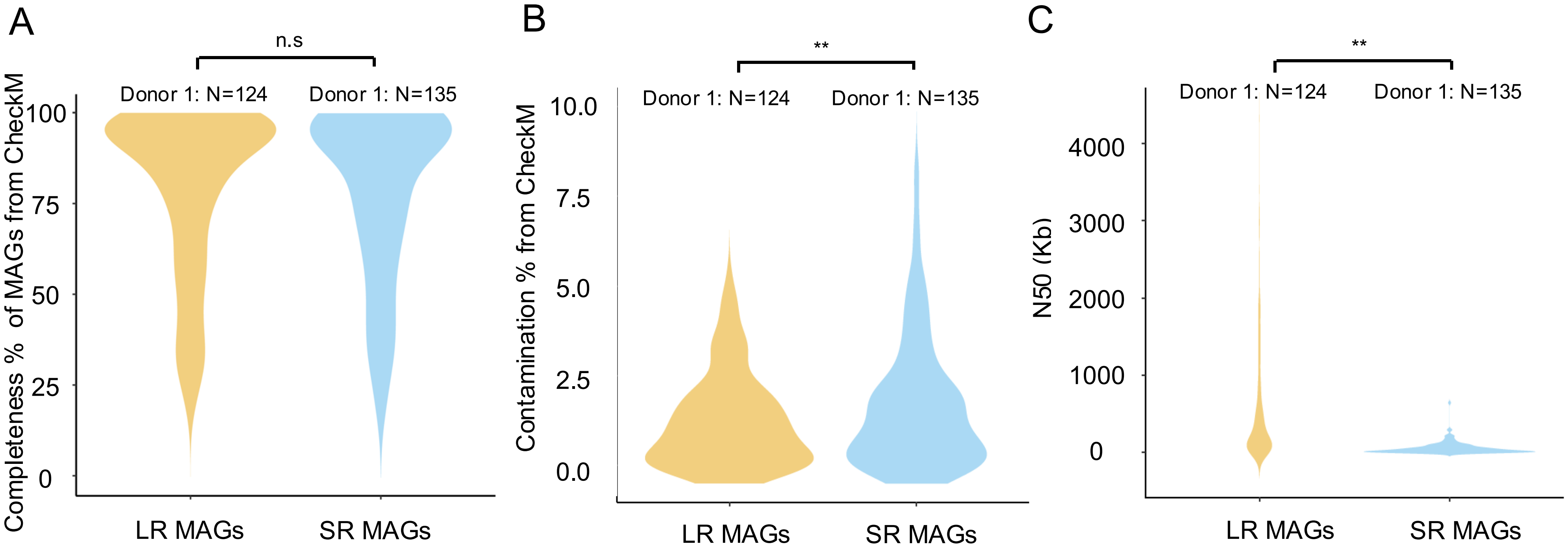
A
n.s
Donor 1: N=124
Donor 1: N=135
Completeness % of MAGs from CheckM
100
75
50
25
0
LR MAGs
SR MAGs
B
**
Donor 1: N=124
Donor 1: N=135
Contamination % from CheckM
10.0
7.5
5.0
2.5
0.0
LR MAGs
SR MAGs
C
**
Donor 1: N=124
Donor 1: N=135
N50 (Kb)
4000
3000
2000
1000
0
LR MAGs
SR MAGs

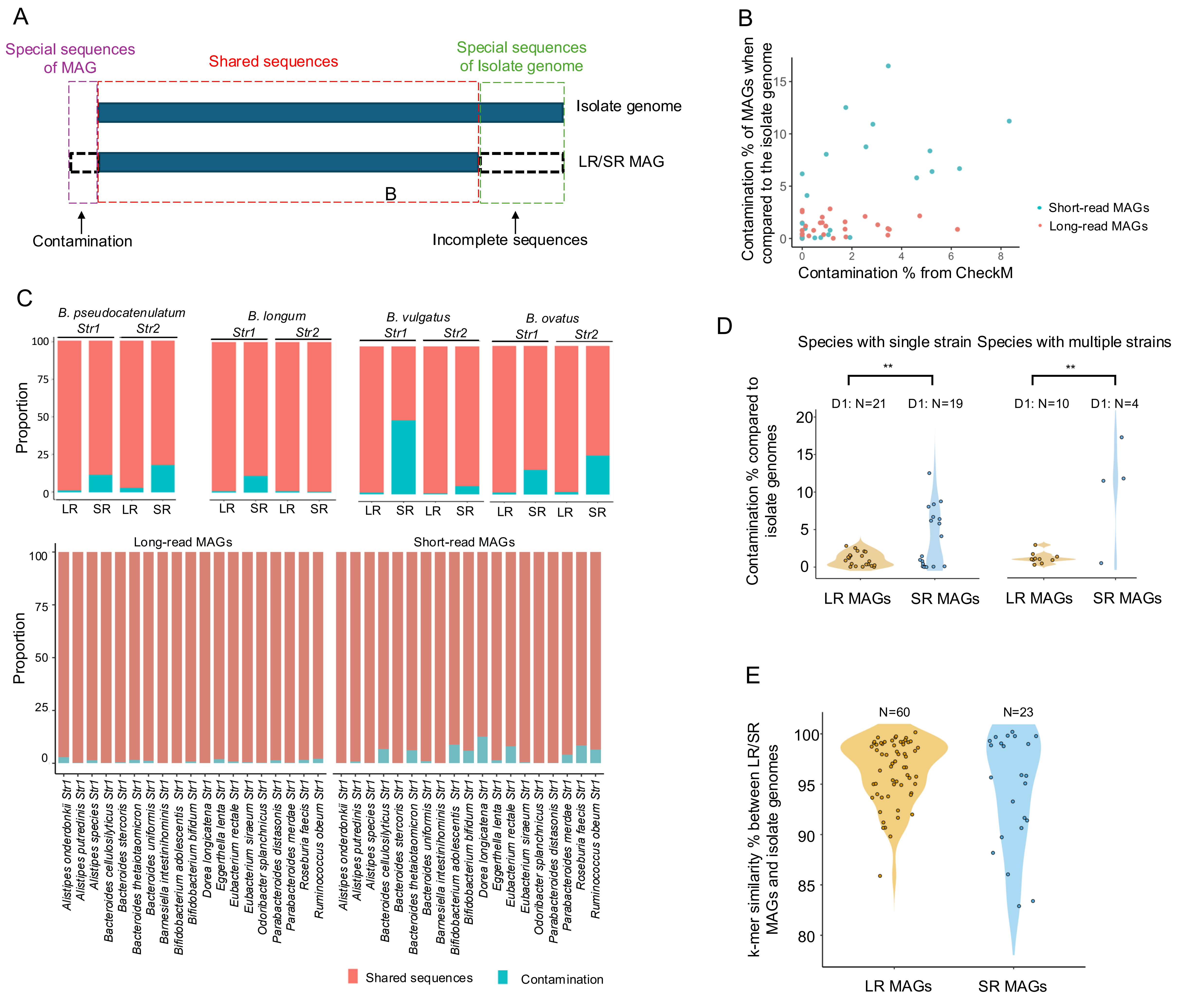

A
Special sequences of MAG
Shared sequences
Special sequences of Isolate genome
Isolate genome
LR/SR MAG
B
Contamination
Incomplete sequences
B
Contamination % of MAGs when compared to the isolate genome
Contamination % from CheckM
Short-read MAGs
Long-read MAGs
C
B. pseudocatenulatum
B. longum
B. vulgatus
B. ovatus
Str1
Str2
Proportion
LR
SR
Long-read MAGs
Short-read MAGs
Alistipes onderdonkii Str1
Alistipes putredinis Str1
Alistipes species Str1
Bacteroides cellulosilyticus Str1
Bacteroides stercoris Str1
Bacteroides thetaiotaomicron Str1
Bacteroides uniformis Str1
Barnesiella intestinihominis Str1
Bifidobacterium adolescentis Str1
Bifidobacterium bifidum Str1
Dorea longicatena Str1
Eggerthella lenta Str1
Eubacterium rectale Str1
Eubacterium siraeum Str1
Odoribacter splanchnicus Str1
Parabacteroides distasonis Str1
Parabacteroides merdae Str1
Roseburia faecis Str1
Ruminococcus obeum Str1
Shared sequences
Contamination
D
Species with single strain
Species with multiple strains
D1: N=21
D1: N=19
D1: N=10
D1: N=4
Contamination % compared to isolate genomes
LR MAGs
SR MAGs
E
N=60
N=23
k-mer similarity % between LR/SR MAGs and isolate genomes
LR MAGs
SR MAGs

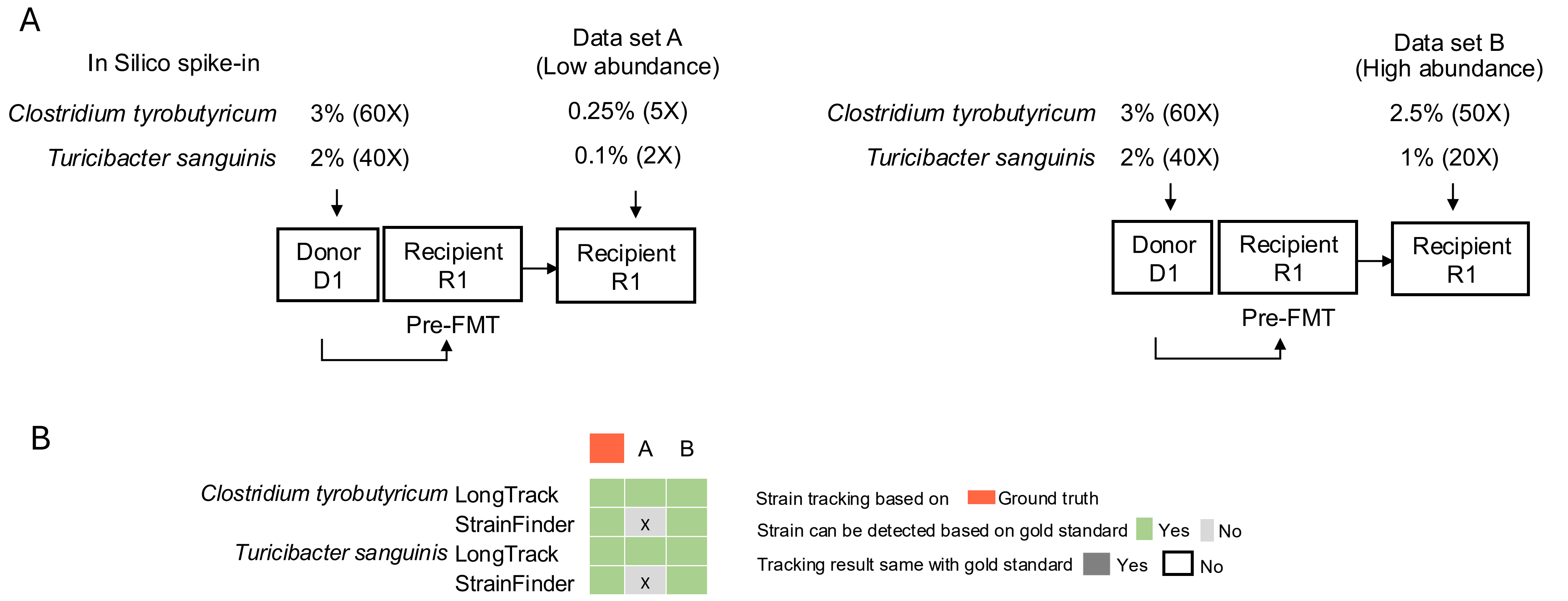
A
In Silico spike-in
Data set A
(Low abundance)
Clostridium tyrobutyricum 3% (60X) 0.25% (5X)
Turicibacter sanguinis 2% (40X) 0.1% (2X)
Donor
D1
Recipient
R1
Recipient
R1
Pre-FMT
Data set B
(High abundance)
Clostridium tyrobutyricum 3% (60X) 2.5% (50X)
Turicibacter sanguinis 2% (40X) 1% (20X)
Donor
D1
Recipient
R1
Recipient
R1
Pre-FMT
B
A
B
Clostridium tyrobutyricum LongTrack
StrainFinder
x
Turicibacter sanguinis LongTrack
StrainFinder
x
Strain tracking based on
Ground truth
Strain can be detected based on gold standard
Yes
No
Tracking result same with gold standard
Yes
No

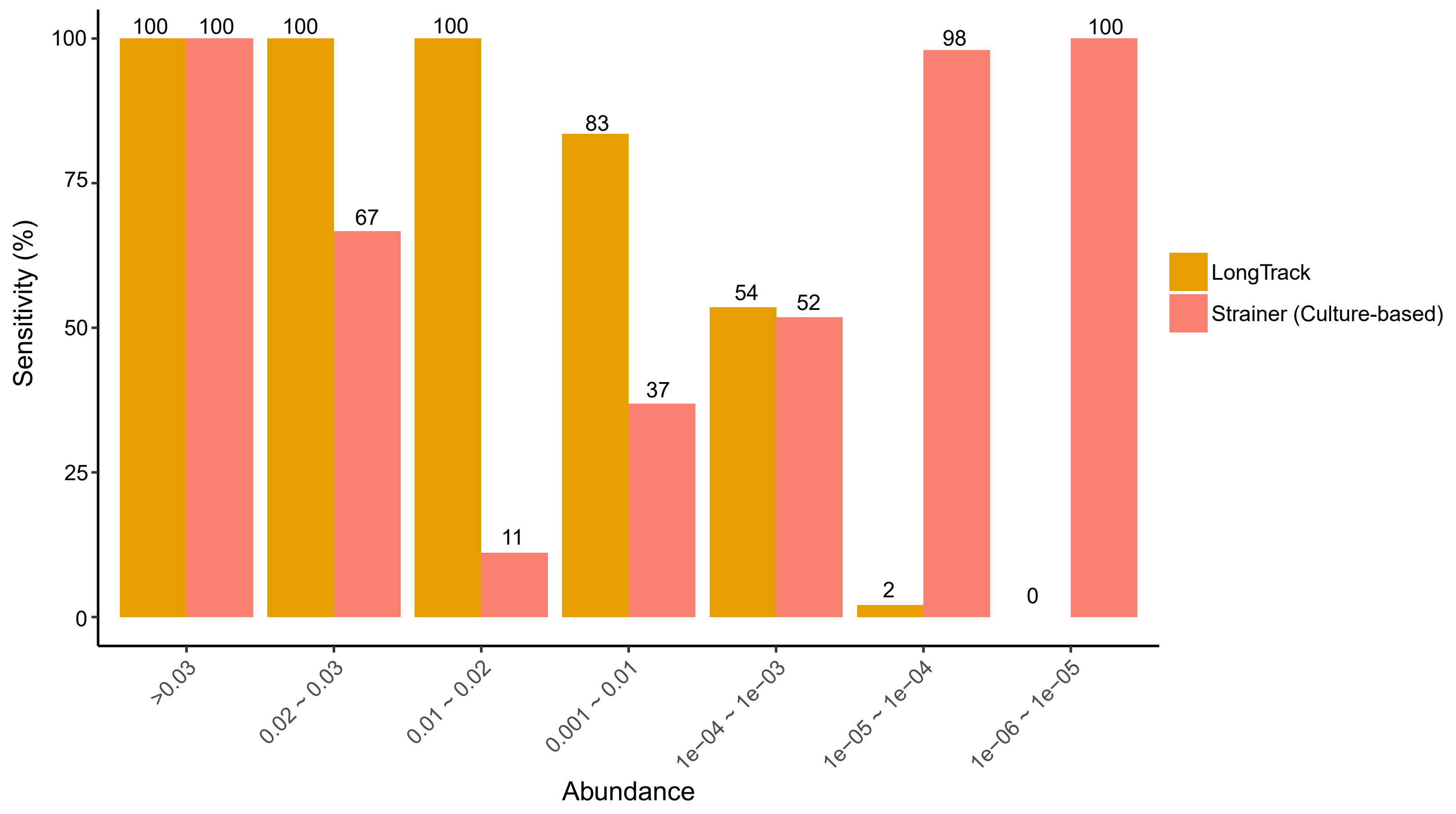

Sensitivity (%)
100
75
50
25
0
100
100
100
67
100
11
83
37
54
52
2
98
0
100
>0.03
0.02 ~ 0.03
0.01 ~ 0.02
0.001 ~ 0.01
1e−04 ~ 1e−03
1e−05 ~ 1e−04
1e−06 ~ 1e−05
Abundance
LongTrack
Strainer (Culture-based)

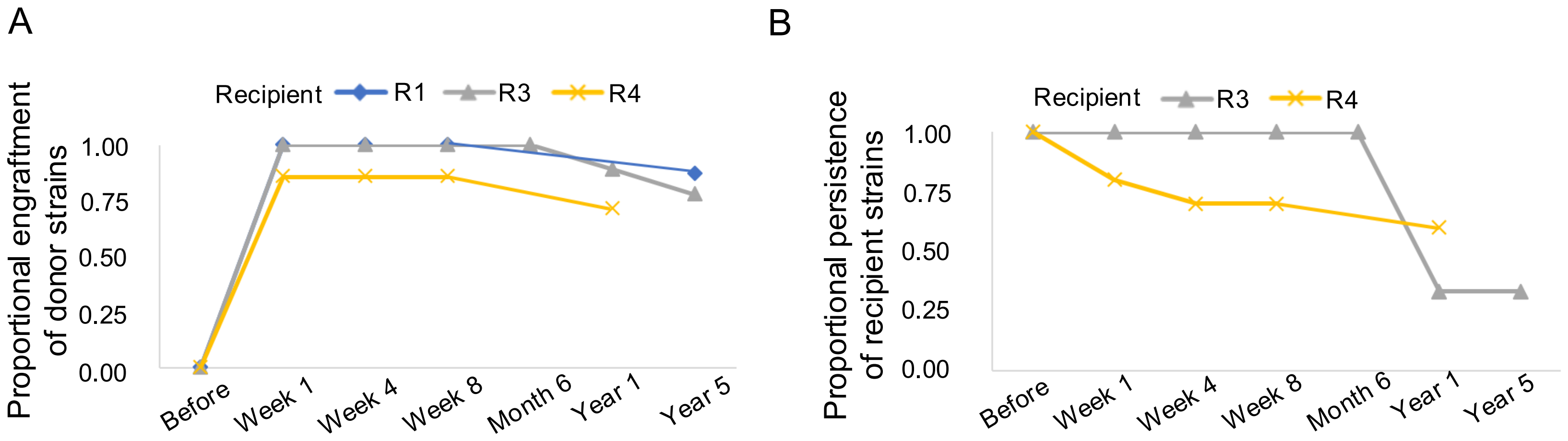
A
Recipient
R1
R3
R4
Proportional engraftment of donor strains
1.00
0.75
0.50
0.25
0.00
Before
Week 1
Week 4
Week 8
Month 6
Year 1
Year 5
B
Recipient
R3
R4
Proportional persistence of recipient strains
1.00
0.75
0.50
0.25
0.00
Before
Week 1
Week 4
Week 8
Month 6
Year 1
Year 5

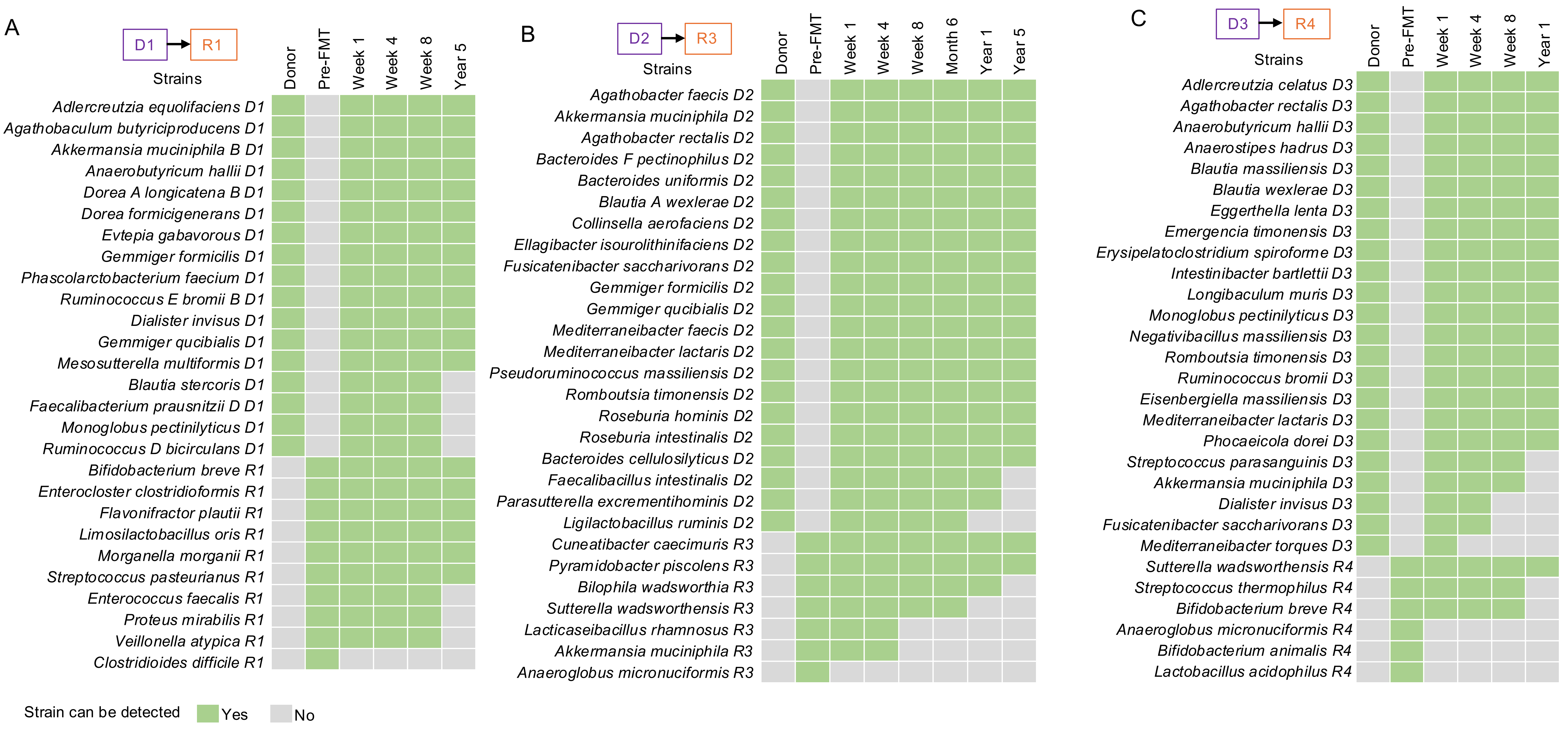
A
D1
R1
Strains
Donor
Pre-FMT
Week 1
Week 4
Week 8
Year 5
Adlercreutzia equolifaciens D1
Agathobaculum butyriciproducens D1
Akkermansia muciniphila B D1
Anaerobutyricum hallii D1
Dorea A longicatena B D1
Dorea formicigenerans D1
Evtepia gabavorous D1
Gemmiger formicilis D1
Phascolarctobacterium faecium D1
Ruminococcus E bromii B D1
Dialister invisus D1
Gemmiger qucibialis D1
Mesosutterella multiformis D1
Blautia stercoris D1
Faecalibacterium prausnitzii D D1
Monoglobus pectinilyticus D1
Ruminococcus D bicirculans D1
Bifidobacterium breve R1
Enterocloster clostridioformis R1
Flavonifractor plautii R1
Limosilactobacillus oris R1
Morganella morganii R1
Streptococcus pasteurianus R1
Enterococcus faecalis R1
Proteus mirabilis R1
Veillonella atypica R1
Clostridioides difficile R1
B
D2
R3
Strains
Donor
Pre-FMT
Week 1
Week 4
Week 8
Month 6
Year 1
Year 5
Agathobacter faecis D2
Akkermansia muciniphila D2
Agathobacter rectalis D2
Bacteroides F pectinophilus D2
Bacteroides uniformis D2
Blautia A wexlerae D2
Collinsella aerofaciens D2
Ellagibacter isourolithinifaciens D2
Fusicatenibacter saccharivorans D2
Gemmiger formicilis D2
Gemmiger qucibialis D2
Mediterraneibacter faecis D2
Mediterraneibacter lactaris D2
Pseudoruminococcus massiliensis D2
Romboutsia timonensis D2
Roseburia hominis D2
Roseburia intestinalis D2
Bacteroides cellulosilyticus D2
Faecalibacillus intestinalis D2
Parasutterella excrementihominis D2
Ligilactobacillus ruminis D2
Cuneatibacter caecimuris R3
Pyramidobacter piscolens R3
Bilophila wadsworthia R3
Sutterella wadsworthensis R3
Lacticaseibacillus rhamnosus R3
Akkermansia muciniphila R3
Anaeroglobus micronuciformis R3
C
D3
R4
Strains
Donor
Pre-FMT
Week 1
Week 4
Week 8
Year 1
Adlercreutzia celatus D3
Agathobacter rectalis D3
Anaerobutyricum hallii D3
Anaerostipes hadrus D3
Blautia massiliensis D3
Blautia wexlerae D3
Eggerthella lenta D3
Emergencia timonensis D3
Erysipelatoclostridium spiroforme D3
Intestinibacter bartlettii D3
Longibaculum muris D3
Monoglobus pectinilyticus D3
Negativibacillus massiliensis D3
Romboutsia timonensis D3
Ruminococcus bromii D3
Eisenbergiella massiliensis D3
Mediterraneibacter lactaris D3
Phocaeicola dorei D3
Streptococcus parasanguinis D3
Akkermansia muciniphila D3
Dialister invisus D3
Fusicatenibacter saccharivorans D3
Mediterraneibacter torques D3
Sutterella wadsworthensis R4
Streptococcus thermophilus R4
Bifidobacterium breve R4
Anaeroglobus micronuciformis R4
Bifidobacterium animalis R4
Lactobacillus acidophilus R4
Strain can be detected
Yes
No

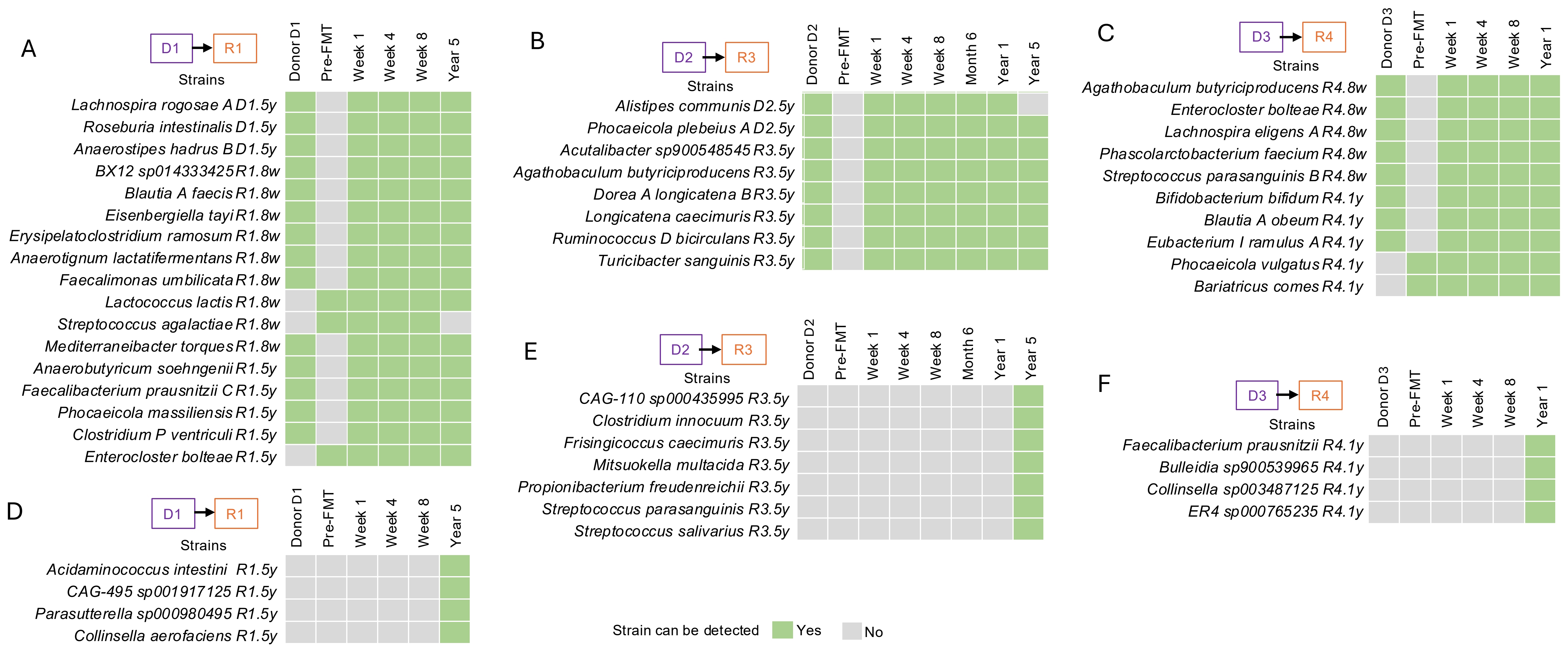

A
D1
R1
Strains
Donor D1
Pre-FMT
Week 1
Week 4
Week 8
Year 5
Lachnospira rogosae A D1.5y
Roseburia intestinalis D1.5y
Anaerostipes hadrus B D1.5y
BX12 sp014333425 R1.8w
Blautia A faecis R1.8w
Eisenbergiella tayi R1.8w
Erysipelatoclostridium ramosum R1.8w
Anaerotignum lactatifermentans R1.8w
Faecalimonas umbilicata R1.8w
Lactococcus lactis R1.8w
Streptococcus agalactiae R1.8w
Mediterraneibacter torques R1.8w
Anaerobutyricum soehngenii R1.5y
Faecalibacterium prausnitzii C R1.5y
Phocaeicola massiliensis R1.5y
Clostridium P ventriculi R1.5y
Enterocloster bolteae R1.5y
B
D2
R3
Strains
Donor D2
Pre-FMT
Week 1
Week 4
Week 8
Month 6
Year 1
Year 5
Alistipes communis D2.5y
Phocaeicola plebeius A D2.5y
Acutalibacter sp900548545 R3.5y
Agathobaculum butyriciproducens R3.5y
Dorea A longicatena B R3.5y
Longicatena caecimuris R3.5y
Ruminococcus D bicirculans R3.5y
Turicibacter sanguinis R3.5y
C
D3
R4
Strains
Donor D3
Pre-FMT
Week 1
Week 4
Week 8
Year 1
Agathobaculum butyriciproducens R4.8w
Enterocloster bolteae R4.8w
Lachnospira eligens A R4.8w
Phascolarctobacterium faecium R4.8w
Streptococcus parasanguinis B R4.8w
Bifidobacterium bifidum R4.1y
Blautia A obeum R4.1y
Eubacterium I ramulus A R4.1y
Phocaeicola vulgatus R4.1y
Bariatricus comes R4.1y
D
D1
R1
Strains
Donor D1
Pre-FMT
Week 1
Week 4
Week 8
Year 5
Acidaminococcus intestini R1.5y
CAG-495 sp001917125 R1.5y
Parasutterella sp000980495 R1.5y
Collinsella aerofaciens R1.5y
E
D2
R3
Strains
Donor D2
Pre-FMT
Week 1
Week 4
Week 8
Month 6
Year 1
Year 5
CAG-110 sp000435995 R3.5y
Clostridium innocuum R3.5y
Frisingicoccus caecimuris R3.5y
Mitsuokella multacida R3.5y
Propionibacterium freudenreichii R3.5y
Streptococcus parasanguinis R3.5y
Streptococcus salivarius R3.5y
F
D3
R4
Strains
Donor D3
Pre-FMT
Week 1
Week 4
Week 8
Year 1
Faecalibacterium prausnitzii R4.1y
Bulleidia sp900539965 R4.1y
Collinsella sp003487125 R4.1y
ER4 sp000765235 R4.1y
Strain can be detected
Yes
No

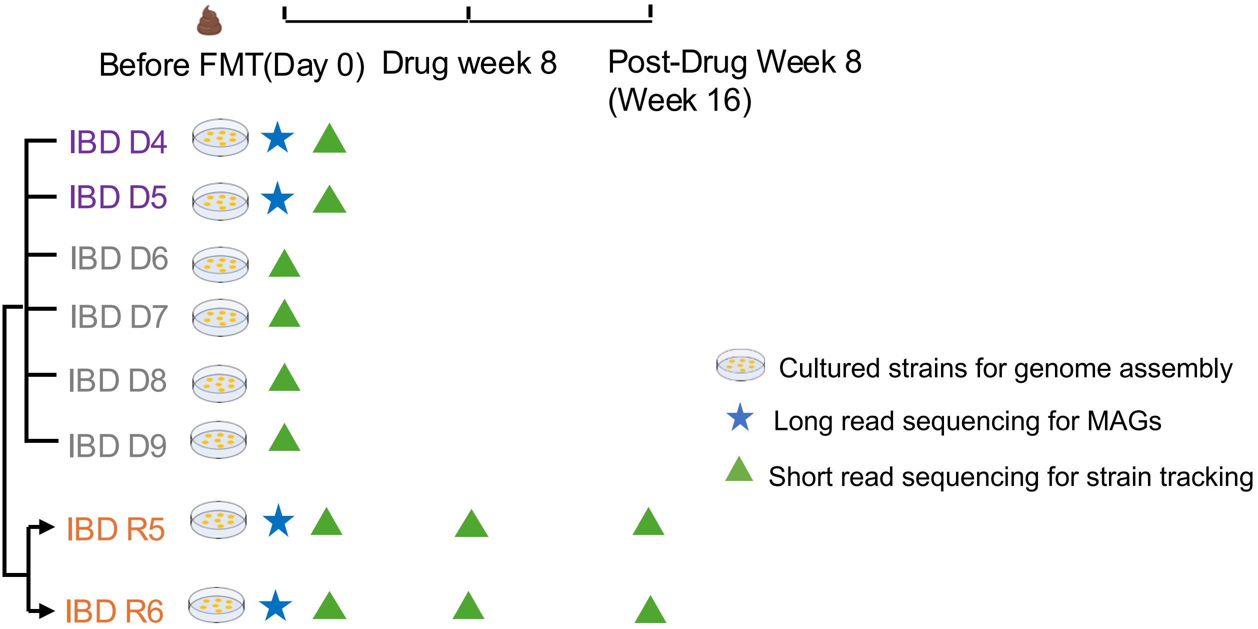


Drug week 8: FMT dosage followed by 40 enemas over 8 weeks, five times per week
Post-Drug Week 8: 8 weeks after cessation of 8 weeks of FMT dosage
