## Supplementary Figures for "Long read metagenomics-based precise tracking of bacterial strains and genomic changes after fecal microbiota transplantation"

### Supplementary Figure

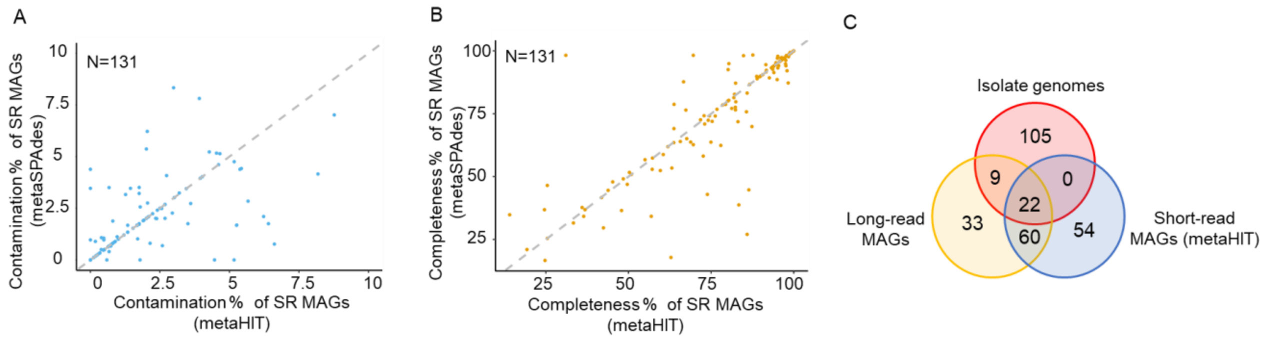


**Supplementary Figure 1. Comparison of meta-assembly methods using metaHIT and metaSPAdes.**

**A.** Contamination levels of 131 shared short-read (SR) MAGs assembled with metaHIT and metaSPAdes are compared, showing no significant differences between the two methods. **B**. Completeness of the same 131 shared SR MAGs assembled with metaHIT and metaSPAdes are also compared, demonstrating no significant differences. **C**. A Venn diagram illustrating the overlap in strain recovery among LR MAGs (yellow), SR MAGs assembled with metaHIT (blue), and isolate genomes (red). LR MAGs recover all 22 strains identified by SR MAGs and include an additional 9 strains not assembled by short-read methods.

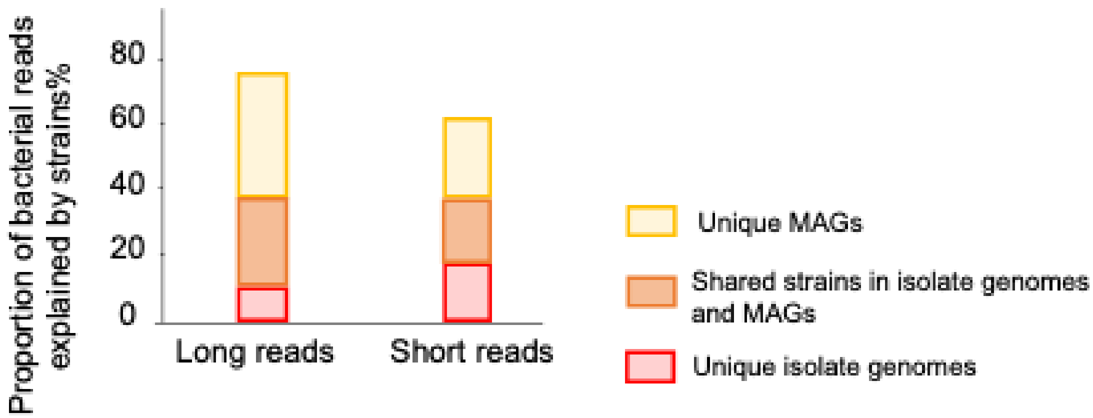


**Supplementary Figure 2.** The proportion of mapped reads from metagenomic sample D1 to the isolate genomes, LR/SR MAGs, or both.

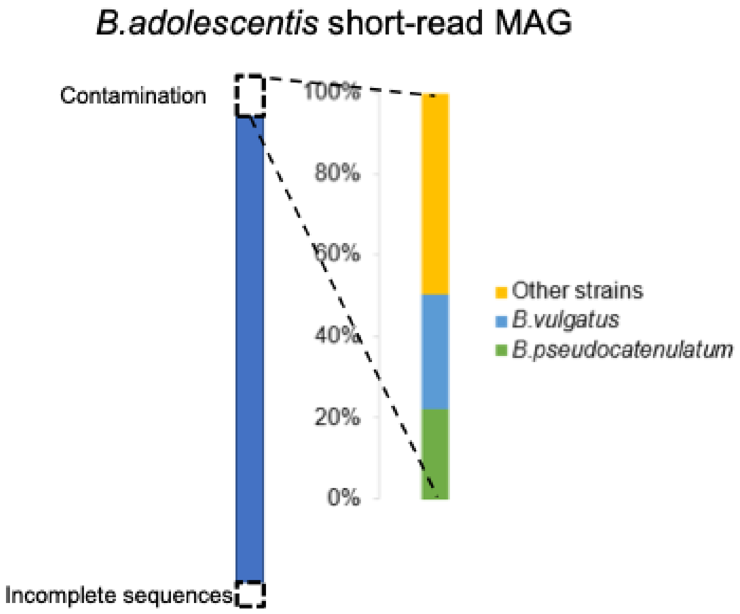


**Supplementary Figure 3.** Contamination composition in *B. adolescentis* short-read MAG. The plot indicates over half of the contamination is collectively attributed to *B. vulgatus* (28%) and *B.* pseudocatenulatum (26%) within the sample donor D1, which led to the subsequent unreliable selection of unique k-mers and the ultimate false positive tracking results in the Year 5 post-FMT sample from R1 (Fig. 3A).

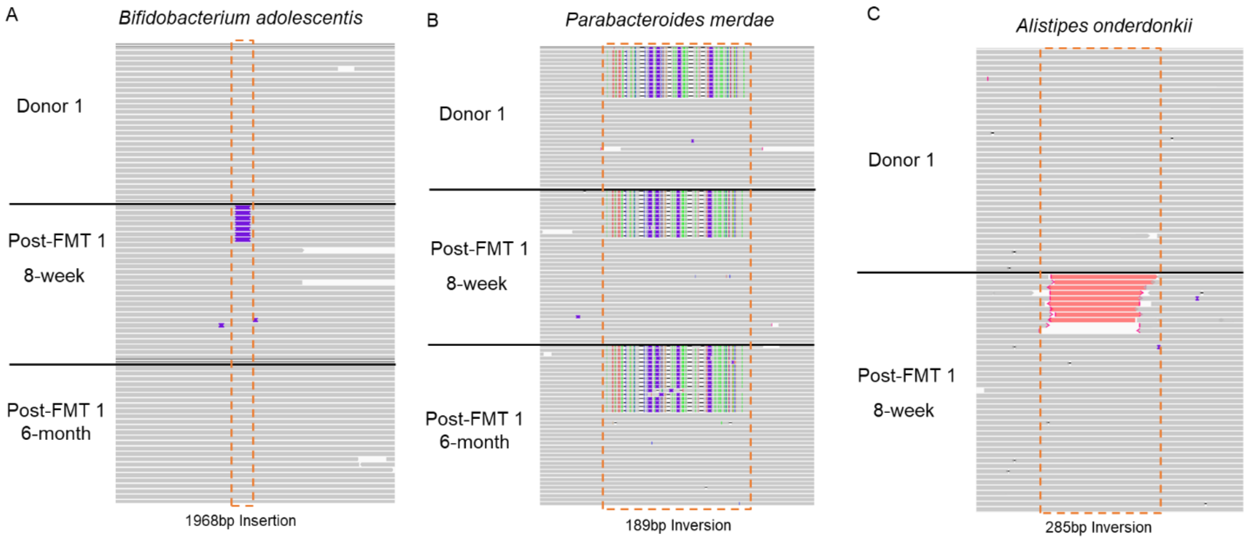


**Supplementary Figure 4. Validation of structural variations using isolate long-read sequencing**

IGV plots demonstrate the validation of structural variations identified through long-read sequencing. **A**, An insertion observed in *Bifidobacterium adolescentis* isolates from Post-FMT at 5 weeks. **B**, An inversion detected in *Parabacteroides merdae* isolates, present in both the donor and Post-FMT samples at 8 weeks and 6 months. **C**, An inversion, highlighted in red within the reads, was identified in *Alistipes onderdonkii* isolates, detected in Post-FMT samples at 8 weeks.
